## Supplemental Information for "Heuristic energy-based cyclic peptide design"

### 1 Cyclic error derivation

Recall that a point with coordinate $\boldsymbol{r_{i+1}}$ in system $i+1$ has coordinate $\boldsymbol{r_i}$ in system $i$, following

$$\boldsymbol{r_i} = \boldsymbol{T_{\theta_{i+1}}} \boldsymbol{R_{\varphi_{i+1}}} \boldsymbol{r_{i+1}} + \boldsymbol{p_i}, \tag{1}$$

where

$$\boldsymbol{T_{\theta_{i+1}}} = \begin{bmatrix} cos(\pi - \theta_{i+1}) & -sin(\pi - \theta_{i+1}) & 0 \\ sin(\pi - \theta_{i+1}) & cos(\pi - \theta_{i+1}) & 0 \\ 0 & 0 & 1 \end{bmatrix},$$

$$\boldsymbol{R_{\varphi_{i+1}}} = \begin{bmatrix} 1 & 0 & 0 \\ 0 & cos(\varphi_{i+1}) & -sin(\varphi_{i+1}) \\ 0 & sin(\varphi_{i+1}) & cos(\varphi_{i+1}) \end{bmatrix}, \boldsymbol{p_i} = \begin{bmatrix} d_i \\ 0 \\ 0 \end{bmatrix}.$$

By choosing system 1 to be at atom $C^\alpha$ of residue $n$, we have bond lengths

$$\begin{aligned} d_1 = d_4 = d_7 = \cdots = d_{3n-2} = d_{3n+1} = d_{C^\alpha} \quad &(\text{bond length } C^\alpha\text{-}C'), \\ d_2 = d_5 = d_8 = \cdots = d_{3n-1} = d_{3n+2} = d_{C'} \quad &(\text{bond length } C'\text{-N}), \\ d_3 = d_6 = d_9 = \cdots = d_{3n} = d_{3n+3} = d_N \quad &(\text{bond length N-}C^\alpha), \end{aligned} \tag{2}$$

bond angles

$$\begin{aligned} \theta_1 = \theta_4 = \theta_7 = \cdots = \theta_{3n-2} = \theta_{3n+1} = \theta_{C^\alpha} \quad &(\text{bond angle N-}C^\alpha\text{-}C'), \\ \theta_2 = \theta_5 = \theta_8 = \cdots = \theta_{3n-1} = \theta_{3n+2} = \theta_{C'} \quad &(\text{bond angle } C^\alpha\text{-}C'\text{-N}), \\ \theta_3 = \theta_6 = \theta_9 = \cdots = \theta_{3n} = \theta_{3n+3} = \theta_N \quad &(\text{bond angle } C'\text{-N-}C^\alpha), \end{aligned} \tag{3}$$

and torsion angles

$$\begin{aligned} &\varphi_1 = \psi_n, \quad \varphi_4 = \psi_1, \quad \varphi_7 = \psi_2, \quad \cdots, \quad \varphi_{3n-2} = \psi_{n-1}, \quad \varphi_{3n+1} = \psi_n, \\ &\varphi_2 = \omega, \quad \varphi_5 = \omega, \quad \varphi_8 = \omega, \quad \cdots, \quad \varphi_{3n-1} = \omega, \quad \varphi_{3n+2} = \omega, \\ &\varphi_3 = \phi_1, \quad \varphi_6 = \phi_2, \quad \varphi_9 = \phi_3, \quad \cdots, \quad \varphi_{3n} = \phi_n, \quad \varphi_{3n+3} = \phi_1, \end{aligned} \tag{4}$$

where $\phi_j$ and $\psi_j$ are torsion angles of residue $j$.

The origin of system $3n+1$ has coordinate $\boldsymbol{r_{3n+1}} = \mathbf{0}$ in system $3n+1$. Using Eq (1), Eq (2), Eq (3), and Eq (4), we have

$$\boldsymbol{r_{3n}} = \begin{bmatrix} d_N \\ 0 \\ 0 \end{bmatrix}, \quad \boldsymbol{r_{3n-1}} = \boldsymbol{T_{\theta_N}} \boldsymbol{R_{\phi_n}} \begin{bmatrix} d_N \\ 0 \\ 0 \end{bmatrix} + \begin{bmatrix} d_{C'} \\ 0 \\ 0 \end{bmatrix} = \boldsymbol{T_{\theta_N}} \begin{bmatrix} d_N \\ 0 \\ 0 \end{bmatrix} + \begin{bmatrix} d_{C'} \\ 0 \\ 0 \end{bmatrix},$$

$$\boldsymbol{r_{3n-2}} = \boldsymbol{T_{\theta_{C'}}} \boldsymbol{R_\omega} \boldsymbol{T_{\theta_N}} \begin{bmatrix} d_N \\ 0 \\ 0 \end{bmatrix} + \boldsymbol{T_{\theta_{C'}}} \begin{bmatrix} d_{C'} \\ 0 \\ 0 \end{bmatrix} + \begin{bmatrix} d_{C^\alpha} \\ 0 \\ 0 \end{bmatrix},$$

$$\boldsymbol{r_{3n-3}} = \boldsymbol{T_{\theta_{C^\alpha}}} \boldsymbol{R_{\psi_{n-1}}} \boldsymbol{T_{\theta_{C'}}} \boldsymbol{R_\omega} \boldsymbol{T_{\theta_N}} \begin{bmatrix} d_N \\ 0 \\ 0 \end{bmatrix} + \boldsymbol{T_{\theta_{C^\alpha}}} \boldsymbol{R_{\psi_{n-1}}} \boldsymbol{T_{\theta_{C'}}} \begin{bmatrix} d_{C'} \\ 0 \\ 0 \end{bmatrix} + \boldsymbol{T_{\theta_{C^\alpha}}} \begin{bmatrix} d_{C^\alpha} \\ 0 \\ 0 \end{bmatrix} + \begin{bmatrix} d_N \\ 0 \\ 0 \end{bmatrix},$$

$$\begin{aligned} \boldsymbol{r_{3n-4}} =& \boldsymbol{T_{\theta_N}} \boldsymbol{R_{\phi_{n-1}}} \boldsymbol{T_{\theta_{C^\alpha}}} \boldsymbol{R_{\psi_{n-1}}} \boldsymbol{T_{\theta_{C'}}} \boldsymbol{R_\omega} \boldsymbol{T_{\theta_N}} \begin{bmatrix} d_N \\ 0 \\ 0 \end{bmatrix} + \boldsymbol{T_{\theta_N}} \boldsymbol{R_{\phi_{n-1}}} \boldsymbol{T_{\theta_{C^\alpha}}} \boldsymbol{R_{\psi_{n-1}}} \boldsymbol{T_{\theta_{C'}}} \begin{bmatrix} d_{C'} \\ 0 \\ 0 \end{bmatrix} \\ &+ \boldsymbol{T_{\theta_N}} \boldsymbol{R_{\phi_{n-1}}} \boldsymbol{T_{\theta_{C^\alpha}}} \begin{bmatrix} d_{C^\alpha} \\ 0 \\ 0 \end{bmatrix} + \boldsymbol{T_{\theta_N}} \begin{bmatrix} d_N \\ 0 \\ 0 \end{bmatrix} + \begin{bmatrix} d_{C'} \\ 0 \\ 0 \end{bmatrix}, \end{aligned}$$

$$\begin{aligned} \boldsymbol{r_{3n-5}} =& \boldsymbol{T_{\theta_{C'}}} \boldsymbol{R_\omega} \boldsymbol{T_{\theta_N}} \boldsymbol{R_{\phi_{n-1}}} \boldsymbol{T_{\theta_{C^\alpha}}} \boldsymbol{R_{\psi_{n-1}}} \boldsymbol{T_{\theta_{C'}}} \boldsymbol{R_\omega} \boldsymbol{T_{\theta_N}} \begin{bmatrix} d_N \\ 0 \\ 0 \end{bmatrix} + \boldsymbol{T_{\theta_{C'}}} \boldsymbol{R_\omega} \boldsymbol{T_{\theta_N}} \boldsymbol{R_{\phi_{n-1}}} \boldsymbol{T_{\theta_{C^\alpha}}} \boldsymbol{R_{\psi_{n-1}}} \boldsymbol{T_{\theta_{C'}}} \begin{bmatrix} d_{C'} \\ 0 \\ 0 \end{bmatrix} \\ &+ \boldsymbol{T_{\theta_{C'}}} \boldsymbol{R_\omega} \boldsymbol{T_{\theta_N}} \boldsymbol{R_{\phi_{n-1}}} \boldsymbol{T_{\theta_{C^\alpha}}} \begin{bmatrix} d_{C^\alpha} \\ 0 \\ 0 \end{bmatrix} + \boldsymbol{T_{\theta_{C'}}} \boldsymbol{R_\omega} \boldsymbol{T_{\theta_N}} \begin{bmatrix} d_N \\ 0 \\ 0 \end{bmatrix} + \boldsymbol{T_{\theta_{C'}}} \begin{bmatrix} d_{C'} \\ 0 \\ 0 \end{bmatrix} + \begin{bmatrix} d_{C^\alpha} \\ 0 \\ 0 \end{bmatrix}. \end{aligned}$$

Let matrix $\boldsymbol{M_i} = \boldsymbol{T_{\theta_{C'}}} \boldsymbol{R_\omega} \boldsymbol{T_{\theta_N}} \boldsymbol{R_{\phi_i}} \boldsymbol{T_{\theta_{C^\alpha}}} \boldsymbol{R_{\psi_i}}$ and $\boldsymbol{q} = \boldsymbol{T_{\theta_{C'}}} \boldsymbol{R_\omega} \boldsymbol{T_{\theta_N}} \begin{bmatrix} d_N \\ 0 \\ 0 \end{bmatrix} + \boldsymbol{T_{\theta_{C'}}} \begin{bmatrix} d_{C'} \\ 0 \\ 0 \end{bmatrix} + \begin{bmatrix} d_{C^\alpha} \\ 0 \\ 0 \end{bmatrix}$, then

$$\boldsymbol{r_{3n-2}} = \boldsymbol{q}, \quad \boldsymbol{r_{3n-5}} = \boldsymbol{M_{n-1}} \boldsymbol{q} + \boldsymbol{q}.$$

If we continue the process, we obtain

$$\begin{aligned} \boldsymbol{r_{3n-8}} &= \boldsymbol{M_{n-2}} \boldsymbol{M_{n-1}} \boldsymbol{q} + \boldsymbol{M_{n-2}} \boldsymbol{q} + \boldsymbol{q}, \\ \boldsymbol{r_{3n-11}} &= \boldsymbol{M_{n-3}} \boldsymbol{M_{n-2}} \boldsymbol{M_{n-1}} \boldsymbol{q} + \boldsymbol{M_{n-3}} \boldsymbol{M_{n-2}} \boldsymbol{q} + \boldsymbol{M_{n-3}} \boldsymbol{q} + \boldsymbol{q}, \quad \cdots \end{aligned}$$

Finally, the requirement for having the same origin is

$$\boldsymbol{r_1} = \boldsymbol{M_1} \boldsymbol{M_2} \cdots \boldsymbol{M_{n-2}} \boldsymbol{M_{n-1}} \boldsymbol{q} + \boldsymbol{M_1} \boldsymbol{M_2} \cdots \boldsymbol{M_{n-2}} \boldsymbol{q} + \cdots + \boldsymbol{M_1} \boldsymbol{M_2} \boldsymbol{q} + \boldsymbol{M_1} \boldsymbol{q} + \boldsymbol{q} = \mathbf{0}. \quad (5)$$

For vectors, the translation $\boldsymbol{p_i}$ in Eq (1) can be ignored. Going through the same derivation process, a vector $\boldsymbol{v_{3n+1}}$ in system $3n+1$ has its vector form

$$\boldsymbol{v_1} = \boldsymbol{M_1 M_2 \cdots M_{n-2} M_{n-1} M_n v_{3n+1}}$$

in system 1. Therefore, the $\boldsymbol{x}$ and $\boldsymbol{y}$ directional vectors in system $3n+1$ have vector forms

$$\boldsymbol{M_1 M_2 \cdots M_{n-2} M_{n-1} M_n e_1}, \quad \boldsymbol{M_1 M_2 \cdots M_{n-2} M_{n-1} M_n e_2},$$

in system 1, respectively, with

$$\boldsymbol{e_1} = \begin{bmatrix} 1 \\ 0 \\ 0 \end{bmatrix}, \quad \boldsymbol{e_2} = \begin{bmatrix} 0 \\ 1 \\ 0 \end{bmatrix}.$$

#### 2 Backbone energy functions

We choose Rosetta's *Ref2015* energy model.[1] We compute energies for the backbone atoms, which are Ramachandran energy, repulsive energy, attractive energy, electrostatic energy, solvation energy, and hydrogen bond energy.

As all backbone residues are assumed to be glycine, we use the permissive, flattened glycine Ramachandran space to calculate backbone Ramachandran energy. We have the Ramachandran energy on the corners of grid cells based on a 10° spacing. For any point inside a grid cell, we calculate its Ramachandran energy by bicubic interpolation of the values on the cell corners. In particular, for a cell with lower left corner at $(\phi_m, \psi_n)$, its bicubic interpolation is

$$E_{rama}^{(m,n)}(\phi, \psi) = \sum_{i=0}^{3} \sum_{j=0}^{3} a_{ij}^{(m,n)} \Big( \frac{\phi - \phi_m}{\Delta\phi} \Big)^i \Big( \frac{\psi - \psi_n}{\Delta\psi} \Big)^j,$$

where $\Delta\phi = \Delta\psi = 10°$. The coefficients $a_{ij}^{(m,n)}$ are set to ensure continuous energy values $E_{rama}$ and partial derivatives $\frac{\partial E_{rama}}{\partial \phi}$, $\frac{\partial E_{rama}}{\partial \psi}$, and $\frac{\partial^2 E_{rama}}{\partial \phi \partial \psi}$ at the four corners. We compute and save these bicubic interpolation coefficients in advance for all cells.

When analyzing backbone atom pair interactions, we follow Rosetta's *Ref2015* energy model.[1] In this model, only atom pairs separated by at least 4 covalent bonds are considered in order to prevent dominance by short-range interactions. We represent backbone atoms as nodes and covalent bonds as edges of length one. By calculating the shortest paths between each atom pair, we compute energies for pairs with path lengths $\geq 4$. This approach aligns with the rationale in the Rosetta paper[1] to "exclude the large repulsive energetic contributions that would otherwise be calculated for atoms separated by fewer than four chemical bonds". For *Van der Waals* interactions, we follow the *Ref2015* model to split the Lennard-Jones 6-12 potential into repulsive and attractive energies. Similarly, we approximate Coulomb's law to compute electrostatic energies, and apply the Lazaridis-Karplus implicit Gaussian exclusion model to compute isotropic solvation energies. Example energy plots are shown in Fig. S1a for the backbone atom pair N and C′.

Hydrogen bond energy consists of three components $E_{hbond}^{HA}$, $E_{hbond}^{AHD}$, and $E_{hbond}^{B_2BAH}$. $H$ refers to the hydrogen atom, and $D$ is its donor (backbone atom N, Fig. S1b). $A$ refers to the acceptor (atom O), $B$ its base (atom C′), and $B2$ is the parent (atom $C^{\alpha}$). The distance between the hydrogen and its acceptor is $d_{HA}$. The bond angle between $A$, $H$, and $D$ is $\theta_{AHD}$. The bond angle between $B$, $A$, and $H$ is $\theta_{BAH}$. The torsion angle between $B2$, $B$, $A$, and $H$ is $\chi_{B2BAH}$. If the bond angles satisy $90° \leq \theta_{AHD}$ and $\theta_{BAH} \leq 180°$, then the energy component $E_{hbond}^{HA}$ forms the curve shown in Fig. S1b; otherwise, $E_{hbond}^{HA} = 0$. Similarly, if $d_{HA} \leq 3.2$ Å and $90° \leq \theta_{BAH} \leq 180°$, then the energy component $E_{hbond}^{AHD}$ forms the curve in Fig. S1b; otherwise, $E_{hbond}^{AHD} = 0$. We draw the Lambert azimuthal project of $E_{hbond}^{B_2BAH}$ in Fig. S1b, with $\theta_{BAH}$ corresponding to the radius and $\chi_{B2BAH}$ rotating counterclockwise. The final hydrogen bond energy $E_{hbond} = w_H w_A f(E_{hbond}^{HA} + E_{hbond}^{AHD} + E_{hbond}^{B_2BAH})$, where $w_H = 1.41$, $w_A = 1.08$ for backbone hydrogen bond, and

$$f(x) = \begin{cases} x & \text{if } x < -0.1 \\ -0.025 + \frac{x}{2} - 2.5x^2 & \text{if } -0.1 \leq x < 0.1 \\ 0 & \text{if } 0.1 \leq x \end{cases}$$

We consider a hydrogen bond to be formed if $E_{hbond} < -0.25$.

Figure S1: **Backbone energy functions.** (A) Example repulsive, attractive, electrostatic, and isotropic solvation energies between backbone atom N and atom C′ are plotted against their distance. (B) The three components of hydrogen bond energy. All energies are in units of kcal/mol.

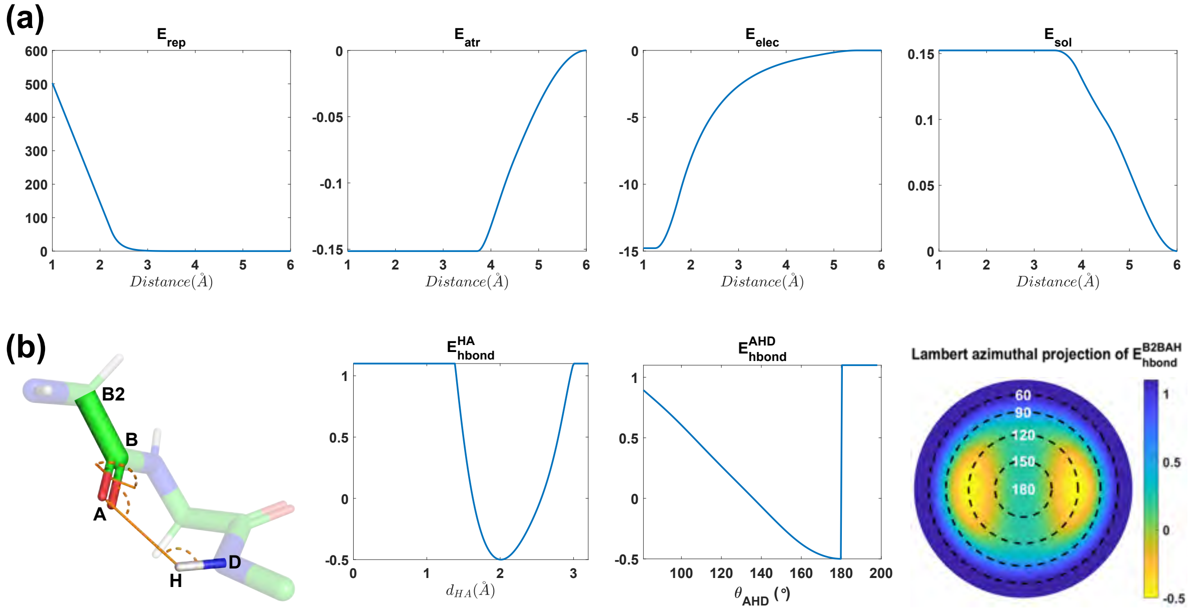

### 3 Layered simulated annealing

To build initial backbone configurations, we partition the glycine Ramachandran map into six torsion bins (Figure S2). For each backbone residue, its initial angles $\phi$ and $\psi$ are chosen randomly from one of the six torsion bin centers. Figure S2 shows an example 7-residue polyglycine chain, with all initial angles chosen as center 1.

Figure S2: **Extended glycine Ramachandran space sampling.** (A) The Ramachandran space is partitioned into six torsion bins, with centers marked and Ramachandran energy (kcal/mol) plotted. A partial simulated annealing path of a residue is drawn for illustration, starting from center 2 with a random move disk of radius $k_0$. At time step $t$, the random move disk shrinks to a radius of $k_t$. (B) An example initial 7-residue polyglycine chain has all torsion angles chosen as center 1. An example good configuration after simulated annealing is labeled with hydrogen bonds and energies. All energies are in units of kcal/mol.

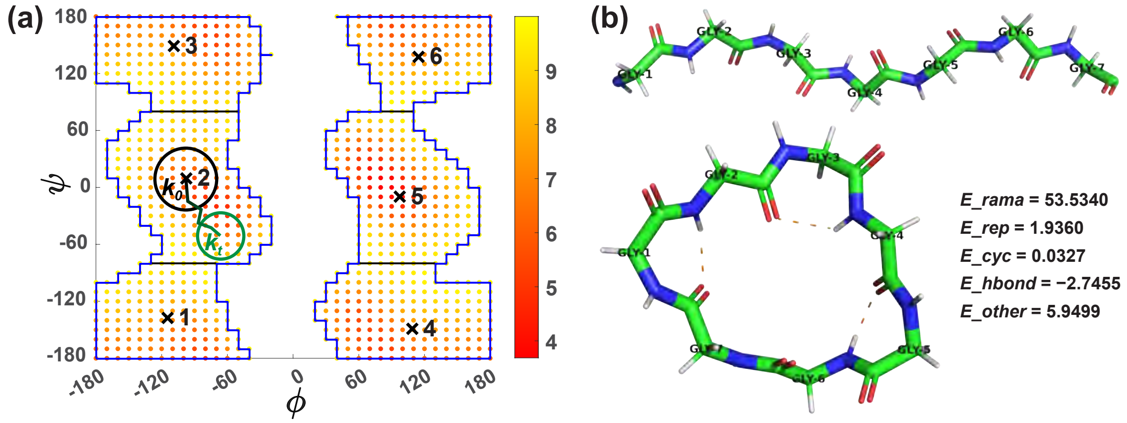


For each initial backbone configuration, we perform the layered simulated annealing. Each layer contains a Metropolis criterion to decide whether the new configuration should be accepted or not. In the cyclic error Metropolis criterion, the energy threshold $E_{thr,cyc}$ is set to be 0.3 for 7 and 15 residues, so that the N-C terminal bond angles and bond lengths deviate little from the ideal values. For 20 and 24 residues, we choose a more lenient $E_{thr,cyc}$ threshold of 1, to find enough backbone candidates under the strict hydrogen bond requirements. Though the N-C terminal bond angles and lengths deviate more, they are still within reasonable ranges to form the peptide bond, and can easily be relaxed to near-ideal geometry by the subsequent application of the Rosetta FastRelax protocol.[2]

For Ramachandran energy, the thresholds are $8n$, where $n$ is the number of residues in the backbone; for hydrogen bonding, we use H-bond count instead of the exact H-bond energy, and the count thresholds are $\lceil n/3 \rceil$ (Table S1). The repulsive and miscellaneous energy thresholds are tuned for a $\sim$15% configuration acceptance rate.

We consider a configuration to be a good backbone candidate, if its repulsive energy $\leq E_{cri,rep}$, cyclic error $\leq E_{cri,cyc}$, and hydsrogen bonds $\geq H_{cri,count}$ (Table S1). We show an example 7-residue

Table S1: **Backbone simulated annealing parameters.** For the four peptide sizes, the thresholds for passing energy tests, the criteria for good backbone candidates, the disk radius parameters for random moves, and the initial temperatures and temperature dropping rates are given.

| $n$ | Energy thresholds | | | | | Candidate criteria | | |
|---|---|---|---|---|---|---|---|---|
| | $E_{thr,rama}$ | $E_{thr,rep}$ | $E_{thr,cyc}$ | $H_{thr,count}$ | $E_{thr,other}$ | $E_{cri,rep}$ | $E_{cri,cyc}$ | $H_{cri,count}$ |
| **7** | 56 | 10 | 0.3 | ≥3 | 10 | 5 | 0.3 | ≥3 |
| **15** | 120 | 15 | 0.3 | ≥5 | – | 10 | 0.3 | ≥5 |
| **20** | 160 | 18 | 1 | ≥7 | – | 13 | 1 | ≥7 |
| **24** | 192 | 20 | 1 | ≥8 | – | 15 | 1 | ≥8 |

| Radius | | Initial temperatures | | | | | Dropping rates | | | | |
|---|---|---|---|---|---|---|---|---|---|---|---|
| $k_0$ | $b$ | $T_{0,rama}$ | $T_{0,rep}$ | $T_{0,cyc}$ | $T_{0,hbond}$ | $T_{0,other}$ | $c_{rama}$ | $c_{rep}$ | $c_{cyc}$ | $c_{hbond}$ | $c_{other}$ |
| 0.9 | 15 | 10 | 20 | 2 | 2 | 20 | 4 | 14 | 16 | 20 | 6 |
| 0.7 | 16 | 20 | 60 | 4 | 4 | – | 2 | 14 | 16 | 18 | – |
| 0.6 | 18 | 25 | 80 | 5 | 5 | – | 4 | 16 | 18 | 22 | – |
| 0.6 | 18 | 30 | 100 | 6 | 6 | – | 4 | 14 | 20 | 22 | – |

good candidate with energies labeled in Fig S2. To design cyclic peptides of intermediate sizes in the future, these thresholds and criteria can be determined using linear interpolation.

A total of $M$=10000 time steps are performed in the layered simulated annealing, and the random move disk radius $k_t$ and temperatures $T_{t,l}$ decrease as a function of time as follows:

$$k_t = \frac{k_0}{1 + b * t/M}, \quad T_{t,l} = \frac{T_{0,l}}{1 + c_l * t/M}.$$

The initial temperature parameters $T_{0,l}$ are chosen to be two times the average energy deviations in test runs, to generate an initial passing probability of $\sim 0.6$ (Table S1). Since these $T_{0,l}$ values exhibit a linear growth pattern, linear interpolation can again be used to determine their values for future cyclic peptide design.

For the other simulated annealing parameters (initial random move disk radius $k_0$, disk shrinking rate $b$, and temperature dropping rates $c_{rama}$, $c_{rep}$, $c_{cyc}$, $c_{hbond}$, and $c_{other}$), approximate ranges are found in test runs and are divided evenly between 3 and 8 values. Within the thousands of possible simulated annealing parameter combinations, we use combinatorial design to select 51 combinations for 7- and 15-residue tests, and 400 combinations for 20- and 24-residue tests (see Combinatorial design).

The pseudocode for our layered simulated annealing algorithm is provided below.

**Algorithm 1** Pseudocode for layered simulated annealing

**Input:** (i) Peptide sequence (all glycine for backbone sampling); (ii) Number of initial points $N_p$; (iii) Simulated annealing parameters $M$, $E_{thr,rama}$, $E_{thr,rep}$, $E_{thr,cyc}$, $H_{thr,count}$, $E_{thr,other}$ (optional), $E_{cri,rep}$, $E_{cri,cyc}$, $H_{cri,count}$, $k_0$, $b$, $T_{0,rama}$, $T_{0,rep}$, $T_{0,cyc}$, $T_{0,hbond}$, $T_{0,other}$ (optional), $c_{rama}$, $c_{rep}$, $c_{cyc}$, $c_{hbond}$, $c_{other}$ (optional).

**Output:** Good backbone candidates and their energies $E_{rama}$, $E_{rep}$, $E_{hbond}$, $E_{other}$ (optional).

```
Randomly select N_p initial points from possible combinations of torsion bin centers
for each initial point angles do
    Calculate initial energies E_rama, E_rep, E_cyc, E_hbond, and E_other, and H-bound count H_count
    N_repeat ← 1, N_backbone ← 0
    while N_repeat ≤ 3 and N_backbone = 0 do
        for time step t from 1 to M do
            k_t ← k_0 / (1+b*t/M)
            Generate a random move within a disk of radius k_t for each residue
            Record the new point angles_new generated

            Calculate Ramachandran energy E_new,rama at the new point        ▷ Rama energy test
            rama_explore ← False, T_t,rama ← T_0,rama / (1+c_rama*t/M)
            if E_new,rama ≤ E_rama or E_new,rama ≤ E_thr,rama then
                rama_explore ← True
            else
                With probability e^((E_rama − E_new,rama)/T_t,rama), set rama_explore True
            end if                                                          ▷ Rama energy test ends

            if rama_explore is True then                                    ▷ Repulsive energy test
                Calculate repulsive energy E_new,rep at the new point
                rep_explore ← False, T_t,rep ← T_0,rep / (1+c_rep*t/M)
                if E_new,rep ≤ E_rep or E_new,rep ≤ E_thr,rep then
                    rep_explore ← True
                else
                    With probability e^((E_rep − E_new,rep)/T_t,rep), set rep_explore True
                end if                               ▷ Metropolis criterion for repulsive energy ends

                if rep_explore is True then                                 ▷ Cyclic error test
                    Calculate cyclic error E_new,cyc at the new point
                    cyc_explore ← False, T_t,cyc ← T_0,cyc / (1+c_cyc*t/M)
```

**if** $E_{new,cyc} \leq E_{cyc}$ **or** $E_{new,cyc} \leq E_{thr,cyc}$ **then**
  *cyc_explore* $\leftarrow$ True
**else**
  With probability $e^{(E_{cyc}-E_{new,cyc})/T_{t,cyc}}$, set *cyc_explore* True
**end if** ▷ Metropolis criterion for cyclic error ends

**if** *cyc_explore* is True **then** ▷ Hydrogen bond energy test
  Calculate hydrogen bond energy $E_{new,hbond}$ at the new point
  *hbond_explore* $\leftarrow$ False, $T_{t,hbond} \leftarrow \frac{T_{0,hbond}}{1+c_{hbond}*t/M}$
  **if** $E_{new,hbond} \leq E_{hbond}$ **or** $H_{new,count} \geq H_{thr,count}$ **then**
    *hbond_explore* $\leftarrow$ True
  **else**
    With probability $e^{(E_{hbond}-E_{new,hbond})/T_{t,hbond}}$, set *hbond_explore* True
  **end if** ▷ Metropolis criterion for hydrogen bond energy ends

  **if** *hbond_explore* is True **then** ▷ Optional miscellaneous energy test
    Calculate miscellaneous energy $E_{new,other}$ at the new point
    *accept* $\leftarrow$ False, $T_{t,other} \leftarrow \frac{T_{0,other}}{1+c_{other}*t/M}$
    **if** $E_{new,other} \leq E_{other}$ **or** $E_{new,other} \leq E_{thr,other}$ **then**
      *accept* $\leftarrow$ True
    **else**
      With probability $e^{(E_{other}-E_{new,other})/T_{t,other}}$, set *accept* True
    **end if** ▷ Metropolis criterion for miscellaneous energy ends

    **if** *accept* is True **then** ▷ Accept the new point
      $angles \leftarrow angles\_new$, $E_{rama} \leftarrow E_{new,rama}$, $E_{rep} \leftarrow E_{new,rep}$,
      $E_{cyc} \leftarrow E_{new,cyc}$, $E_{hbond} \leftarrow E_{new,hbond}$, $H_{count} \leftarrow H_{new,count}$,
      $E_{other} \leftarrow E_{new,other}$
      **if** $E_{rep} \leq E_{cri,rep}$ **and** $E_{cyc} \leq E_{cri,cyc}$ **and** $H_{count} \geq H_{cri,count}$ **then**
        Record *angles* and its energies as a good backbone candidate
        $N_{backbone} \leftarrow N_{backbone} + 1$
      **end if** ▷ The backbone is recorded as a candidate
    **end if** ▷ The new point is accepted
  **end if** ▷ Miscellaneous energy test ends
**end if** ▷ Hydrogen bond energy test ends
**end if** ▷ Cyclic error test ends
**end if** ▷ Repulsive energy test ends
**end for**
$N_{repeat} \leftarrow N_{repeat} + 1$
**end while**
**end for**

### 4 Combinatorial design

The goal of combinatorial design is to obtain well-spaced random samples from the cross-product of parameter-values.[3] In our case, we are performing a simulated annealing parameter choice, as described in Layered simulated annealing. Specifically, our combinatorial design method guarantees that for any pair of parameters P1 and P2, every possible combination of the discrete values of P1 and P2, such as P1.v1 and P2.v2, is present in at least one initial parameter setting. Note that this differs from exhaustive search because, for example, P1.v1 and P2.v2 may be in the same initial setting as some values P3.v3 and P4.v4, but never in the same initial setting as some other values P3.v3' and P4.v4'.

While an exhaustive search would require 61440 parameter settings for 7 residue backbone sampling (7 parameters $k_0$, $b$, $c_{rama}$, $c_{rep}$, $c_{cyc}$, $c_{hbond}$, and $c_{other}$, each between 3 and 8 values), this form of combinatorial design requires only 51 settings. The same settings are used for the 15 residue simulated annealing parameter optimization.

For 20 and 24 residues, we employ a special form of combinatorial design that has pivots. It uses the generic combinatorial design to find settings of the non-pivot parameters, and repeats the settings for every value of the pivot parameters. We choose initial random move disk radius $k_0$ and disk shrinking rate $b$ as pivots, because, in test runs, they have a large impact on successfully finding good backbones that satisfy the repulsive energy, cyclic error, and hydrogen bond requirements. Parameter $k_0$ has four discrete values (0.6, 0.7, 0.8, and 0.9), $b$ has five values (15, 16, 17, 18, and 19), and the other parameters yield 20 settings by the generic combinatorial design. As a result, the pivot combinatorial design generates 400 ($20 \times 4 \times 5$) parameter settings.

For each parameter combination, after setting the initial angle as center 1 for all residues (Fig 3a of main text), we repeat the layered simulated annealing 20 times for 7 and 15 residues, and 10 times for 20 and 24 residues. In each run of simulated annealing, we record the number of good candidate backbones produced (satisfying criteria in Table S1). The optimal parameter combination is the one which produces candidate backbones in more than half of the simulated annealing runs and has the most candidates in total. For future design of other intermediate sizes, similar parameter optimization can be performed, and we suggest parameter ranges of 0.6-0.9 for $k_0$, 15-19 for $b$, 2-4 for $c_{rama}$, 12-18 for $c_{rep}$, 14-20 for $c_{cyc}$, 16-22 for $c_{hbond}$, and 4-10 for $c_{other}$.

Simulations were performed on the Greene supercomputer clusters at the New York University's High Performance Computing facilities. Each compute node in the Greene clusters has two 24-core Intel Cascade Lake Platinum 8268 chips and 192 GB memory. The parameter combination selection process took 9.6, 16, 33.6, and 41.6 CPU hours for 7, 15, 20, and 24 residues, respectively.

### 5 Torsion angle FastRelax script

```
<ROSETTASCRIPTS>
```

```
    # The SCOREFXNS section defines scoring functions
    <SCOREFXNS>
            # The current Rosetta default scorefunction
            <ScoreFunction name="ref" weights="ref2015" />
            # Use chainbreak to maintain cyclic
            <ScoreFunction name="ref_chainbreak" weights="ref2015" >
                    <Reweight scoretype="chainbreak" weight="15.0" />
            </ScoreFunction>
    </SCOREFXNS>
    # The SIMPLE_METRICS section allows users to configure metrics used to measure
properties of a structure.
    <SIMPLE_METRICS>
            # Metric to measure backbone hydrogen bonds
            <PeptideInternalHbondsMetric name="internal_hbonds" />
            # Metric to measure score/energy
            <TotalEnergyMetric name="score" scorefxn="ref" />
    </SIMPLE_METRICS>
    # The FILTERS section allows users to configure filters. These measure
properties of a structure and make decisions, based on the measured properties,
about whether to discard the current structure.
    <FILTERS>
            # Filter to avoid score function artifact of having more than two
hydrogen bonds to carbonyls
            <OversaturatedHbondAcceptorFilter name="oversat" scorefxn="ref"
max_allowed_oversaturated="0" consider_mainchain_only="false"/>
            # Filter to ensure a minimum number of hbonds (peptide size n/3)
            <PeptideInternalHbondsFilter name="min_internal_hbonds" hbond_cutoff="7"
/>
    </FILTERS>
    # The MOVERS section allows users to define movers, which are Rosetta modules
that modify a structure in some way.
    <MOVERS>
            # A mover to declare a bond connecting the termini (i.e. to cyclize the
peptide). Note that the user needs to input variable %%Nres%% (peptide size) when
running this relaxation script. Use option "-parser:script_vars Nres=20" for 20
residue relaxation.
            <DeclareBond name="peptide_bond1" res1="1" atom1="N" atom2="C"
res2="%%Nres%%" add_termini="true" />
            # Three repeats of fastrelax
            <FastRelax name="frlx" scorefxn="ref_chainbreak" repeats="3"
ramp_down_constraints="false" >
                    # All side-chain and backbone torsion angles can move
                    <MoveMap name="frlx_mm" >
                            <Chain number="1" chi="true" bb="true" />
                    </MoveMap>
```

```
            </FastRelax>
            # These movers allow the simple metrics to be run
            <RunSimpleMetrics name="measure_internal_hbonds"
    metrics="internal_hbonds" />
            <RunSimpleMetrics name="measure_score" metrics="score" />
        </MOVERS>
        # The PROTOCOLS section is the section in which the user invokes the modules
    defined above in linear sequence to define a protocol:
        <PROTOCOLS>
            <Add mover="peptide_bond1" />
            <Add mover="frlx" />
            # A side-effect of the DeclareBond mover is the correction of positions
    of H and O atoms that depend on the peptide bond. We re-invoke it here for that
    purpose.
            <Add mover="peptide_bond1" />
            <Add filter="min_internal_hbonds" />
            <Add filter="oversat" />
        </PROTOCOLS>
        # The OUTPUT section allows the user to define output settings. Here, we specify
    the scoring function that will be used to score the output structure.
        <OUTPUT scorefxn="ref"/>
</ROSETTASCRIPTS>
```

### 6 Small macrocycle FastDesign script

```
<ROSETTASCRIPTS>
        <SCOREFXNS>
            <ScoreFunction name="ref" weights="ref2015" />
            # The default scorefunction with increased hydrogen bond weights, and
    with aa_composition, aspartimide_penalty, and chainbreak scores activated.
            <ScoreFunction name="ref_highhbond" weights="ref2015" >
                <Reweight scoretype="hbond_lr_bb" weight="5.0" />
                <Reweight scoretype="hbond_sr_bb" weight="5.0" />
                <Reweight scoretype="aa_composition" weight="1.0" />
                <Reweight scoretype="aspartimide_penalty" weight="1.0" />
                <Reweight scoretype="chainbreak" weight="15.0" />
            </ScoreFunction>
        </SCOREFXNS>
        # The PACKER_PALETTES section defines the residues available for design.
        <PACKER_PALETTES>
            <CustomBaseTypePackerPalette name="palette" additional_residue_types
    ="DALA,DASP,DGLU,DPHE,DHIS,DILE,DLYS,DLEU,DMET,DASN,DPRO,DGLN,DARG,DSER,DTHR,
    DVAL,DTRP,DTYR" />
        </PACKER_PALETTES>
```

```
    # The RESIDUE_SELECTORS section allows users to select residues.
    <RESIDUE_SELECTORS>
            # Select residues with mainchain phi torsion values greater than zero.
These positions will be restricted to becoming D-amino acids during design.
            <Phi name="posPhi" select_positive_phi="true" />
            # Select residues with mainchain phi torsion values less than zero.
These positions will be restricted to becoming L-amino acids during design.
            <Phi name="negPhi" select_positive_phi="false" />
    </RESIDUE_SELECTORS>
    <SIMPLE_METRICS>
            <PeptideInternalHbondsMetric name="internal_hbonds" />
    </SIMPLE_METRICS>
    <FILTERS>
            <OversaturatedHbondAcceptorFilter name="oversat" scorefxn="ref"
max_allowed_oversaturated="0" consider_mainchain_only="false"/>
            <PeptideInternalHbondsFilter name="min_internal_hbonds" hbond_cutoff="5"
/>
    </FILTERS>
    # The TASKOPERATIONS section allows users to control side-chain identity.
    <TASKOPERATIONS>
            # Task operation to read a resfile defining the D-amino acids, used for
design at positions with mainchain phi torsion values greater than zero.
            <ReadResfile name="d_res" filename="d_res.txt" selector="posPhi"/>
            # Task operation to read a resfile defining the L-amino acids, used for
design at positions with mainchain phi torsion values less than zero.
            <ReadResfile name="l_res" filename="l_res.txt" selector="negPhi"/>
    </TASKOPERATIONS>
    <MOVERS>
            <DeclareBond name="peptide_bond1" res1="1" atom1="N" atom2="C"
res2="%%Nres%%" add_termini="true" />
            # Composition constraints add a nonlinearly-ramping penalty for
deviation from a desired amino acid composition written in the .comp file.
            <AddCompositionConstraintMover name="addcompcsts"
filename="desired_makeup.comp" />
            # The FastDesign mover performs alternating rounds of sequence design
and torsion-space energy minimization, while ramping the repulsive term in the
scorefunction (fa_rep).
            <FastDesign name="fdes" scorefxn="ref_highhbond" repeats="3"
task_operations="d_res,l_res" packer_palette="palette" ramp_down_constraints="false"
>
                    <MoveMap name="fdes_mm" >
                            <Chain number="1" chi="true" bb="true" />
                    </MoveMap>
            </FastDesign>
            <RunSimpleMetrics name="measure_internal_hbonds"
```

```
    metrics="internal_hbonds" />
        </MOVERS>
        <PROTOCOLS>
                <Add mover="peptide_bond1" />
                <Add mover="addcompcsts" />
                <Add mover="fdes" />
                <Add mover="peptide_bond1" />
                <Add filter="oversat" />
                <Add filter="min_internal_hbonds" />
        </PROTOCOLS>
        <OUTPUT scorefxn="ref"/>
</ROSETTASCRIPTS>
```

#### l_res.txt

```
PIKAA ADEFHIKLMNPQRSTVWY
start
```

#### d_res.txt

```
PIKAA X[DALA]X[DASP]X[DGLU]X[DPHE]X[DHIS]X[DILE]X[DLYS]X[DLEU]X[DMET]X[DASN]X[DPRO]
X[DGLN]X[DARG]X[DSER]X[DTHR]X[DVAL]X[DTRP]X[DTYR]
```

#### desired_makeup.comp

```
# At least two proline residues. These can be L- or D- (or mixed).
PENALTY_DEFINITION
TYPE PRO DPR
DELTA_START -2
DELTA_END 1
PENALTIES 500 10 0 0
ABSOLUTE 2
BEFORE_FUNCTION QUADRATIC
AFTER_FUNCTION CONSTANT
END_PENALTY_DEFINITION

# At least one L-asp or L-glu.
PENALTY_DEFINITION
TYPE ASP GLU
DELTA_START -1
DELTA_END 1
PENALTIES 200 0 0
ABSOLUTE 1
```

```
BEFORE_FUNCTION QUADRATIC
AFTER_FUNCTION CONSTANT
END_PENALTY_DEFINITION

# At least one positively-charged residue
PENALTY_DEFINITION
TYPE LYS ARG DLY DAR
DELTA_START -1
DELTA_END 1
PENALTIES 200 0 0
ABSOLUTE 1
BEFORE_FUNCTION QUADRATIC
AFTER_FUNCTION CONSTANT
END_PENALTY_DEFINITION
```

### 7 Large macrocycle FastDesign script

```
<ROSETTASCRIPTS>
        <SCOREFXNS>
                <ScoreFunction name="ref" weights="ref2015" />
                <ScoreFunction name="ref_highhbond" weights="ref2015" >
                        <Reweight scoretype="hbond_lr_bb" weight="10.0" />
                        <Reweight scoretype="hbond_sr_bb" weight="10.0" />
                        <Reweight scoretype="aa_composition" weight="1.0" />
                        <Reweight scoretype="aspartimide_penalty" weight="1.0" />
                        <Reweight scoretype="chainbreak" weight="25.0" />
                </ScoreFunction>
        </SCOREFXNS>
        <PACKER_PALETTES>
                <CustomBaseTypePackerPalette name="palette" additional_residue_types
    ="DALA,DASP,DGLU,DPHE,DHIS,DILE,DLYS,DLEU,DMET,DASN,DPRO,DGLN,DARG,DSER,DTHR,
    DVAL,DTRP,DTYR" />
        </PACKER_PALETTES>
        <RESIDUE_SELECTORS>
                <Phi name="posPhi" select_positive_phi="true" />
                <Phi name="negPhi" select_positive_phi="false" />
                # Select the most buried residues
                <Layer name="select_core" select_core="true" select_boundary="false"
    select_surface="false" core_cutoff="2.5" surface_cutoff="1.0" />
                # Select partially buried residues
                <Layer name="select_boundary" select_core="false" select_boundary="true"
    select_surface="false" core_cutoff="2.5" surface_cutoff="1.0" />
                # Select fully solvent-exposed residues
                <Layer name="select_surface" select_core="false" select_boundary="false"
    select_surface="true" core_cutoff="2.5" surface_cutoff="1.0" />
```

```
    </RESIDUE_SELECTORS>
    <SIMPLE_METRICS>
            <PeptideInternalHbondsMetric name="internal_hbonds" />
    </SIMPLE_METRICS>
    <FILTERS>
            <OversaturatedHbondAcceptorFilter name="oversat" scorefxn="ref"
max_allowed_oversaturated="0" consider_mainchain_only="false"/>
            <PeptideInternalHbondsFilter name="min_internal_hbonds" hbond_cutoff="7"
/>
    </FILTERS>
    <TASKOPERATIONS>
            <ReadResfile name="d_res" filename="d_res.txt" selector="posPhi"/>
            <ReadResfile name="l_res" filename="l_res.txt" selector="negPhi"/>
            # At buried positions, restrict to PFAMILYVW and D-aa equivalents
            <RestrictToSpecifiedBaseResidueTypes name="core_restrictions"
base_types="PRO,PHE,ALA,MET,ILE,LEU,TYR,VAL,TRP,DPRO,DPHE,DALA,DMET,DILE,DLEU,
DTYR,DVAL,DTRP" selector="select_core"/>
            # At semi-buried positions, PAILYVNQDERKSTH and D-aa equivalents
            <RestrictToSpecifiedBaseResidueTypes name="boundary_restrictions"
base_types="PRO,ALA,ILE,LEU,TYR,VAL,ASN,GLN,ASP,GLU,ARG,LYS,SER,THR,HIS,DPRO,
DALA,DILE,DLEU,DTYR,DVAL,DASN,DGLN,DASP,DGLU,DARG,DLYS,DSER,DTHR,DHIS"
selector="select_boundary"/>
            # At surface-exposed positions, PANQDERKSTH and D-aa equivalents
            <RestrictToSpecifiedBaseResidueTypes name="surf_restrictions"
base_types="PRO,ALA,ASN,GLN,ASP,GLU,ARG,LYS,SER,THR,HIS,DPRO,DALA,DASN,DGLN,
DASP,DGLU,DARG,DLYS,DSER,DTHR,DHIS" selector="select_surface"/>
    </TASKOPERATIONS>
    <MOVERS>
            <DeclareBond name="peptide_bond1" res1="1" atom1="N" atom2="C"
res2="%%Nres%%" add_termini="true" />
            <AddCompositionConstraintMover name="addcompcsts"
filename="desired_makeup.comp" />
            <FastDesign name="fdes" scorefxn="ref_highhbond" repeats="3"
task_operations="d_res, l_res, core_restrictions, boundary_restrictions,
surf_restrictions" packer_palette="palette" ramp_down_constraints="false" >
                    <MoveMap name="fdes_mm" >
                            <Chain number="1" chi="true" bb="true" />
                    </MoveMap>
            </FastDesign>
            <RunSimpleMetrics name="measure_internal_hbonds"
metrics="internal_hbonds" />
    </MOVERS>
    <PROTOCOLS>
            <Add mover="peptide_bond1" />
            <Add mover="addcompcsts" />
```

```
                <Add mover="fdes" />
                <Add mover="peptide_bond1" />
                <Add filter="oversat" />
                <Add filter="min_internal_hbonds" />
        </PROTOCOLS>
        <OUTPUT scorefxn="ref"/>
</ROSETTASCRIPTS>
```

#### 8 ClusterGen stability analysis

For macrocycles having 15-24 residues, the Ramachandran-stability filtering method fails to find sufficient alternative backbone conformations due to the exponentially larger search space. There are six different types of Ramachandran spaces (see Fig. S3). The L amino acid Ramachandran spaces hardly overlap with those of the D amino acids, so the chance that a residue's torsion angles fit into the corresponding Ramachandran space is roughly 1/2. This gives a probability of $\sim \frac{1}{2^{15}/15}$ or $\frac{1}{2^{24}/24}$ of finding a compatible backbone candidate with a 15- or 24-residue sequence, respectively. Therefore, efficient search of low-energy alternative conformations is needed.

To form the initial population of ClusterGen, we perform two separate layered simulated annealing, one for low energy, and one for low RMSD. The low energy simulated annealing is the same as in Layered simulated annealing, while the low RMSD simulated annealing has only the cyclic error test and a RMSD test (see pseudocode below). All parameters follow Table 1 of main text, and the newly added RMSD parameters are threshold $E_{thr,rmsd}$=1.5 Å, the 15- and 20-residue criterion $E_{cri,rmsd}$=1.5 Å, the 24-residue criterion $E_{cri,rmsd}$=1.7 Å, the initial temperature $T_{0,rmsd}$=1, and the temperature dropping rate $c_{rmsd}$=50, chosen by test runs. For both low energy and low RMSD simulated annealing procedures of 15 residues, we randomly select 1000 initial points from possible combinations of torsion bin centers corresponding to the designed sequence (Fig. S3). For 20 residues, we select 10000 initial points for each SA procedure. For 24 residues, we select 20000 initial points for low energy SA, and 40000 for low RMSD SA.

The sampled backbones are then subject to FastRelax. We sort the relaxed structures in ascending order of their energies and initiate energy-based clustering. Each time, we select the lowest-energy structure from the remaining unclustered structures (the first one in the sorted list) to serve as a new cluster center. Then, we measure the backbone-heavy-atom RMSDs of the other unclustered structures from this new center. We assign those having RMSDs smaller than 0.5 Å to this cluster, and remove them from the sorted list. We repeat this process until all structures have been assigned to clusters. Note that we record only coordinates of backbone heavy atoms during the genetic algorithm to save memory and computation time, but we relax entire structures including sidechains during FastRelax to obtain full energies.

In the genetic algorithm, each generation undergoes crossover, mutation, and selection (see pseudocode below). Crossover involves checking if a pair of parents can exchange residues within a

Figure S3: **Ramachandran spaces for L and D amino acids.** The six different dypes of Ramachandran spaces correspond to L- and D-proline, L- and D-valine (also isoleucine and threonine), and L- and D-alanine (also remaining amino acids). The Ramachandran space centers are marked and the Ramachandran energies are in units of kcal/mol.

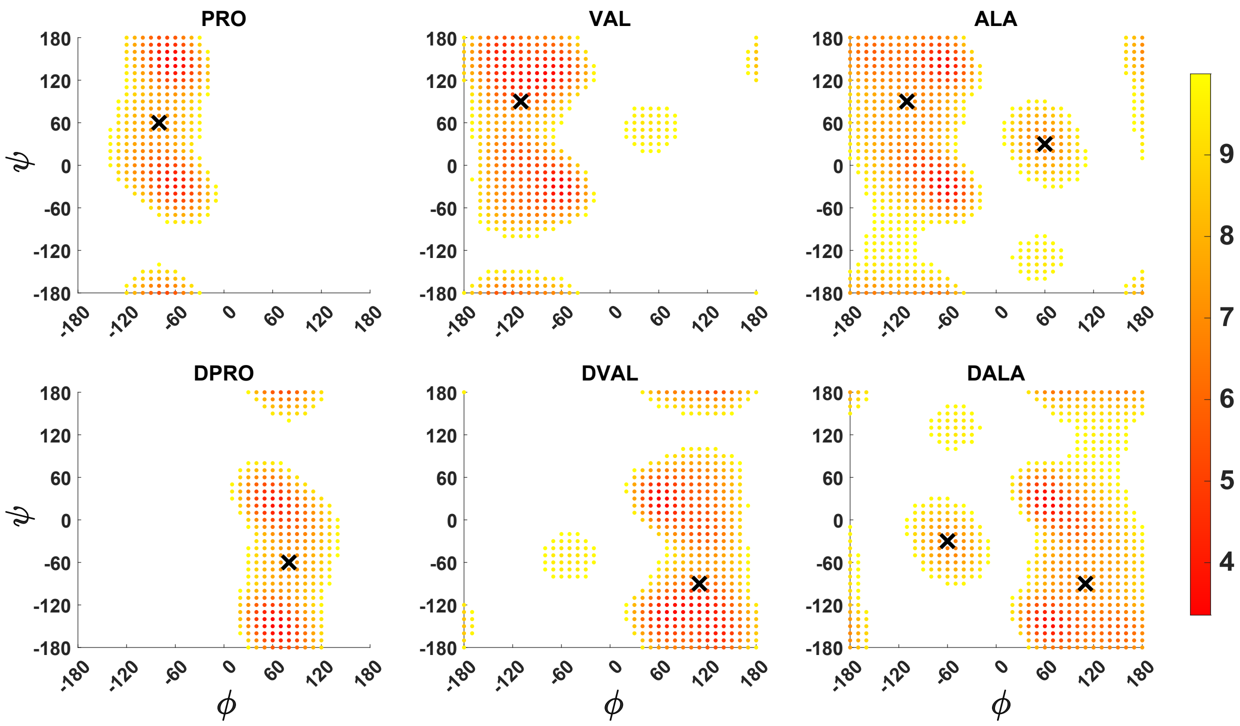


designated region. More specifically, for two breakpoint residues $b1$ and $b2$ (i.e., exchange residues $b1 + 1$ to $b2$), we use the Kabsch algorithm[4] to align eight atoms of the two parents: $C^\alpha$ and $C'$ of residues $b1$ and $b2$, atoms N and $C^\alpha$ of residues $b1 + 1$ and $b2 + 1$ (see Fig. S4). The algorithm first translates both sets of atoms to have their centroids at the origin. Then, it computes the covariance matrix between the two sets of atoms and performs singular value decomposition on this matrix to determine the optimal rotation matrix. This rotation minimizes the sum of the squared distances between corresponding atoms, and the root mean of this sum is the RMSD. If the alignment yields an RMSD $\leq rsmd_{xover}$ (0.5 Å) per atom, the parents exchange these residues to generate two children.

Given a mutation region spanning residues $d1$ to $d2$, we add a random perturbation of up to $p_{max} = 10°$ to the torsion angles of residue $d1$. Subsequently, we adjust the torsion angles of the remaining residues to ensure a smooth connection with residue $d2 + 1$ (see Fig. S4). Specifically, we employ a simulated annealing approach with $M_{mut} = 1000$ steps. In each step $t$, we add new perturbations of $\leq p_t$ to the current torsion angles of residues $d1 + 1$ to $d2$, where $p_t = \frac{10}{1+9*t/1000}$. If a perturbation steps out of the associated Ramachandran space, it is discarded. For the new

Figure S4: **ClusterGen crossover and mutation.** (A) Example crossover of two 15-residue backbones. (B) Example mutation of a 15-residue backbone.

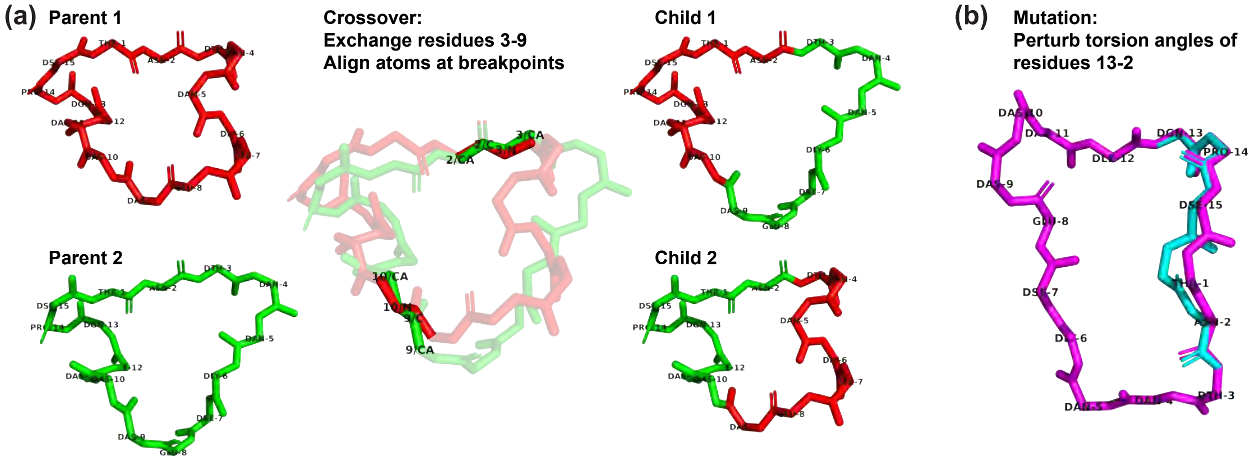


torsion angles, we calculate the coordinates of atoms N and $C^{\alpha}$ of residue $d2+1$, and sum their distances from the original N and $C^{\alpha}$ positions. If the resulting distance sum, denoted $D_{new}$, is smaller than the current sum $D$, the new torsion angles are accepted. Otherwise, the new angles still have a probability of $e^{(D-D_{new})/T_{t,D}}$ to be accepted, where $T_{t,D} = \frac{T_{0,D}}{1+c_D*t/1000}$, and $T_{0,D} = 1$ and $c_D = 20$ based on test runs. The simulated annealing process stops whenever $D \leq D_{mut}$ (0.3 Å), indicating a smooth connection between the mutated residues and residue $d2$+1.

The crossover and mutation regions are randomly chosen from any section in the length range of $L_{xover}$ and $L_{mut}$ (3-8 for 15 residue macrocycles, 3-10 for 20 residues, and 4-12 for 24 residues). We randomly sort all possible parent pairs, and continue crossover until reaching $s_{thr,xover} = 1.5 * N_{GA}$ children. Similarly, for mutation, we randomly sample 100*$N_{GA}$ backbones from the current population, and continue mutation until reaching $s_{thr,mut} = 1.5 * N_{GA}$ mutants.

ClusterGen stability analyzes were performed on the Greene supercomputer clusters at the New York University's High Performance Computing facilities. In Fig. S5, we plot the computation time spent on different components of ClusterGen, including the initial-population-generating layered simulated annealing, the Matlab part of the genetic algorithm (crossover, mutation, and selection), and the FastRelax part of the genetic algorithm (Cartesian relaxation after crossover and mutation), assuming 96 cores used. The increase in time spent on simulated annealing is due to the expanded initial points. In the genetic algorithm, about 70% of the computation time is spent on Cartesian relaxation.

Figure S5: **ClusterGen computation time breakdown.** For macrocycle designs of 15, 20, and 24 residues, we breakdown the computation CPU hours per ClusterGen stability analysis, using NYU Greene HPC.

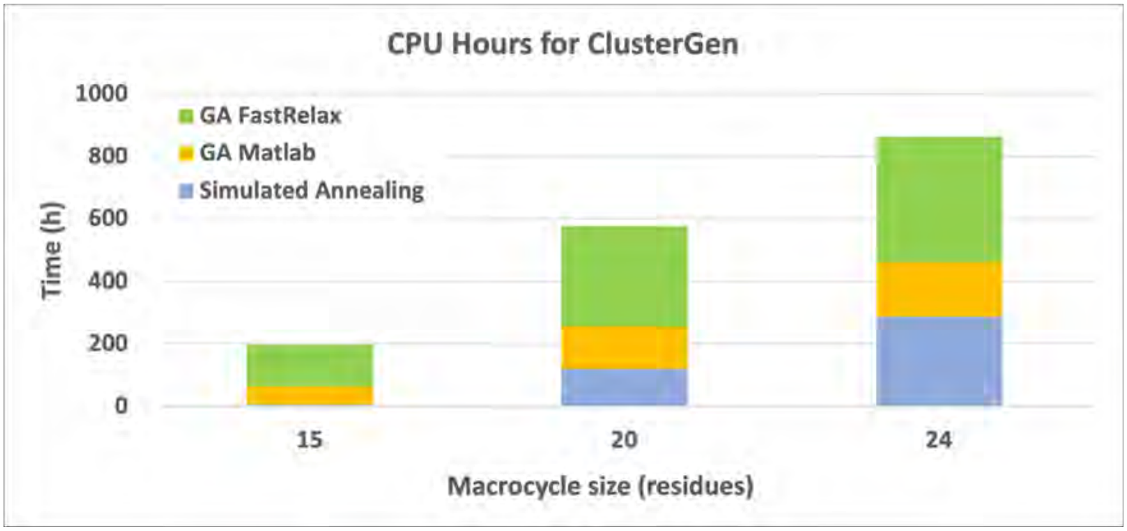

**Algorithm 2** Pseudocode for ClusterGen

**Step 1. Generating initial population by simulated annealing**

**Input:** (i) Designed sequence and structure; (ii) Number of simulated annealing initial points for low energy $N_{p,energy}$ and for low RMSD $N_{p,rmsd}$; (iii) Simulated annealing parameters $M$, $E_{thr,rama}$, $E_{thr,rep}$, $E_{thr,cyc}$, $E_{thr,rmsd}$, $H_{thr,count}$, $E_{cri,rep}$, $E_{cri,cyc}$, $E_{cri,rmsd}$, $H_{cri,count}$, $k_0$, $b$, $T_{0,rama}$, $T_{0,rep}$, $T_{0,cyc}$, $T_{0,hbond}$, $T_{0,rmsd}$, $c_{rama}$, $c_{rep}$, $c_{cyc}$, $c_{hbond}$, $c_{rmsd}$.

**Output:** Alternative backbones for the designed sequence.

---

Randomly select $N_{p,energy}$ initial points from possible combinations of torsion bin centers (Fig. S3) corresponding to the designed sequence ▷ Low energy SA starts
**for** each initial point **do**
  Perform the layered simulated annealing with Ramachandran energy test, repulsive energy test, cyclic error test, and hydrogen bond energy test as in Algorithm 1
  Record points that satisfy the repulsive energy, cyclic error, and hydrogen bond count criteria as alternative backbones
**end for** ▷ Low energy SA ends

Randomly select $N_{p,rmsd}$ initial points ▷ Low RMSD SA starts
**for** each initial point *angles* **do**
  Calculate initial $E_{cyc}$ and backbone-heavy-atom RMSD ($E_{rmsd}$) from the designed structure
  **for** time step $t$ from 1 to $M$ **do**
    Generate a random move for each residue within a disk of radius $\frac{k_0}{1+b*t/M}$
    Record the new point *angles_new* generated
    Perform cyclic error test, with parameters $T_{0,cyc}, c_{cyc}, E_{thr,cyc}$
    **if** cyclic error test passed **then** ▷ RMSD test
      Calculate RMSD $E_{new,rmsd}$ at the new point
      $accept \leftarrow$ False, $T_{t,rmsd} \leftarrow \frac{T_{0,rmsd}}{1+c_{rmsd}*t/M}$
      **if** $E_{new,rmsd} \leq E_{rmsd}$ **or** $E_{new,rmsd} \leq E_{thr,rmsd}$ **then**
        $accept \leftarrow$ True
      **else**
        With probability $e^{(E_{rmsd}-E_{new,rmsd})/T_{t,rmsd}}$, set *accept* True
      **end if** ▷ Metropolis criterion for RMSD ends

      **if** *accept* is True **then** ▷ Accept the new point
        $angles \leftarrow angles\_new$, $E_{cyc} \leftarrow E_{new,cyc}$, $E_{rmsd} \leftarrow E_{new,rmsd}$
        **if** $E_{cyc} \leq E_{cri,cyc}$ **and** $E_{rmsd} \leq E_{cri,rmsd}$ **then**
          Record *angles* as an alternative backbone
        **end if**
      **end if** ▷ The new point is accepted
    **end if** ▷ RMSD test ends
  **end for**
**end for** ▷ Low RMSD SA ends

**Step 2. Torsion angle relaxation (section 5) for alternative backbones**

**Input:** Alternative backbones for the designed sequence.

**Output:** Relaxed alternative structures with sidechains added and energies computed.

---

**Step 3. Energy-based clustering**

**Input:** (i) Relaxed alternative structures with energies; (ii) Genetic algorithm population $N_{GA}$; (iii) Clustering RMSD cutoff $rmsd_{cutoff}$ (0.5 Å).

**Output:** $2 * N_{GA}$ lowest-energy cluster centers.

---

Sort the $N$ relaxed structures in ascending order of their energies
$library \leftarrow [1, 2, 3, ..., N]$, $centers \leftarrow [\quad]$
**while** *library* not empty **do**
  $center \leftarrow library(1)$, $members \leftarrow [\quad]$
  **for** each element $e$ in *library* **do**
    **if** backbone RMSD between $e$ and $center < rmsd_{cutoff}$ **then**
      add $e$ to *members*
    **end if**
    remove *members* from *library*, add *center* to *centers*
  **end for**
**end while**
Record the $2 * N_{GA}$ lowest-energy cluster centers

---

**Step 4. Genetic algorithm**

**Input:** (i) Initial population of $2 * N_{GA}$ lowest-energy cluster centers; (ii) Population size $N_{GA}$, number of generations $M_{GA}$, and population decrement $R_{GA}$; (iii) Crossover parameters: $L_{xover}$, $s_{thr,xover}$, $rmsd_{xover}$; (iv) Mutation parameters: $L_{mut}$, $s_{thr,mut}$, $D_{mut}$, $p_{max}$, $M_{mut}$, $T_{0,D}$, $C_D$.

**Output:** Alternative structures with energies for the energy landscape.

---

**for** generation from 1 to $M_{GA}$ **do**
  Crossover children size $s_{xover} \leftarrow 0$ ▷ Crossover starts
  **for** each pair of two parent backbones in the population **do**
    Randomly select a crossover length from the range $L_{xover}$
    Randomly select breakpoint residue $b1$, and add length to find breakpoint $b2$

    Obtain the following atoms' coordinates for both parents:
      $C^{\alpha}$ and $C'$ atoms in residues $b1$ and $b2$
      $N$ and $C^{\alpha}$ atoms in residues $b1 + 1$ and $b2 + 1$

    Align these atoms from the two parents, and calculate RMSD
    **if** RMSD $\leq rmsd_{xover}$ **then**
      Exchange residues $b1 + 1$ to $b2$ of the two parents (Fig. S4)
      Record the two crossover children, $s_{xover} \leftarrow s_{xover} + 2$
      Stop the crossover for loop if $s_{xover} \geq s_{thr,xover}$
    **end if**
  **end for** ▷ Crossover ends

Randomly sample $100 * N_{GA}$ backbone from the population ▷ Mutation starts

Mutation children size $s_{mut} \leftarrow 0$

**for** each sample **do**

Randomly select a mutation length from the range $L_{mut}$

Randomly select the starting residue $d1$, and add length to find the ending residue $d2$

Add random angle perturbations $\leq p_{max}$ to $\phi, \psi$ of residue $d1$

Calculate new positions of atoms $N$ and $C^{\alpha}$ of residue $d2+1$

Sum distances $D$ between these atoms' new positions and their original positions

Step index $t \leftarrow 1$ ▷ Mutation simulated annealing starts

**while** $D > D_{mut}$ and $t \leq M_{mut}$ **do**

Generate random perturbations $\leq \frac{p_{max}}{1+(p_{max}-1)*t/M_{mut}}$ for $\phi, \psi$ of residues $d1+1$ to $d2$

Set the $\phi, \psi$ perturbations of a residue to 0 if it leaves the corresponding Ramachandran space

Calculate new positions of atoms $N$ and $C^{\alpha}$ of residue $d2+1$

Sum distances $D_{new}$ between these atoms' new positions and their original positions

$T_D \leftarrow \frac{T_{0,D}}{(1+c_D*t/M_{mut})}$ ▷ Metropolis criterion starts

**if** $D_{new} \leq D$ **then**

Add the random perturbations, $D \leftarrow D_{new}$

**else**

With probability $e^{(D-D_{new})/T_D}$, add the random perturbations and $D \leftarrow D_{new}$

**end if** ▷ Metropolis criterion ends

$t \leftarrow t+1$

**end while** ▷ Mutation simulated annealing ends

**if** $D \leq D_{mut}$ **then**

Mutate atom positions in residues $d1$ to $d2$ induced by the angle perturbations

Record the mutant child, $s_{mut} \leftarrow s_{mut}+1$

Stop the mutation for loop if $s_{mut} \geq s_{thr,mut}$

**end if**

**end for** ▷ Mutation ends

Perform FastRelax Cartesian relaxation (section 9) for the crossover and mutant children

Perform energy-based clustering (Step 3)

Select $N_{GA}$ lowest-energy cluster centers as the next generation

Record cluster centers with energies $< 0$ as alternative structures in the energy landscape

$N_{GA} \leftarrow N_{GA} - R_{GA}$

**end for**

### 9 Cartesian coordinate FastRelax script

```
<ROSETTASCRIPTS>
	<SCOREFXNS>
		<ScoreFunction name="ref" weights="ref2015" />
		# For Cartesian relaxation
		<ScoreFunction name="ref_cartesian" weights="ref2015" >
			<Reweight scoretype="chainbreak" weight="15.0" />
			<Reweight scoretype="cart_bonded" weight="0.5" />
			<Reweight scoretype="pro_close" weight="0.0" />
		</ScoreFunction>
	</SCOREFXNS>
	<SIMPLE_METRICS>
		<PeptideInternalHbondsMetric name="internal_hbonds" />
		<TotalEnergyMetric name="score" scorefxn="ref" />
		# Metric to measure RMSD from designed structure to produce energy
  landscape
		<RMSDMetric name="rmsd_ref" use_native="true" super="true"
  rmsd_type="rmsd_protein_bb_heavy_including_O" />
	</SIMPLE_METRICS>
	<FILTERS>
		<OversaturatedHbondAcceptorFilter name="oversat" scorefxn="ref"
  max_allowed_oversaturated="0" consider_mainchain_only="false"/>
		# Only 2 minimum hbonds are required for energy landscape sampling
		<PeptideInternalHbondsFilter name="min_internal_hbonds" hbond_cutoff="2"
  />
	</FILTERS>
	<MOVERS>
		<DeclareBond name="peptide_bond1" res1="1" atom1="N" atom2="C"
  res2="%%Nres%%" add_termini="true" />
		# A Cartesian relaxation is performed, trying to adjust the bond angles
  and lengths to their ideal values.
		<FastRelax name="frlx_Cartesian" scorefxn="ref_cartesian" repeats="3"
  ramp_down_constraints="false" cartesian="true" bondangle="true" bondlength="true">
			<MoveMap name="frlx_mm" >
				<Chain number="1" chi="true" bb="true" />
			</MoveMap>
		</FastRelax>
		<RunSimpleMetrics name="measure_internal_hbonds"
  metrics="internal_hbonds" />
		<RunSimpleMetrics name="measure_score" metrics="score" />
		<RunSimpleMetrics name="measure_rmsd" metrics="rmsd_ref" />
	</MOVERS>
	<PROTOCOLS>
		<Add mover="peptide_bond1" />
```

```
                <Add mover="frlx_Cartesian" />
                <Add mover="peptide_bond1" />
                <Add filter="min_internal_hbonds" />
                <Add filter="oversat" />
                <Add mover="measure_rmsd" />
        </PROTOCOLS>
        <OUTPUT scorefxn="ref"/>
</ROSETTASCRIPTS>
```

#### 10 MD and REMD protocol

Our molecular dynamics (MD) simulation protocol started with a rapid energy minimization, which used the Particle Mesh Ewald method[5] for long-range electrostatic interactions with a cutoff of 1 nm, and constrained the lengths of all bonds that involved a hydrogen atom. The water molecules were rigid. A leapfrog Verlet integrator was used with timestep of 1 fs.

After energy minimization, we conducted a 100 ps constant volume and temperature (NVT) ensemble equilibration using Langevin middle integrator, which does the LFMiddle discretization.[6] The temperature was set to be 300 K, the collision rate 1/ps, and the timestep 2 fs. Finally, we ran a 1 $\mu$s constant pressure and temperature (NPT) ensemble using the same Langevin middle integrator setting, together with a Monte Carlo Barostat[7,8] at pressure 1 bar. We wrote trajectory frames every 10 ps. For trajectory analysis and RMSD calculations, wed use the Python package *MDAnalysis*.[9]

For replica exchange molecular dynamics (REMD) simulations, the initial preparation involved solvation, energy minimization, and equilibration within an NVT ensemble, following the same protocols as described previously. The prepared system was then replicated across a series of temperatures ranging from 300 K to 500 K. We chose the temperatures using a webserver[10] to have a predicted exchange probability of 0.25.

A constant pressure of 1 bar was maintained for all replicas. During each iteration, 1000 steps of Langevin dynamics integration were performed with collision rate 1/ps and timestep 2 fs. At the end of the iteration, replicas with neighboring temperatures had a chance to swap, following the Metropolis criterion.[11] We ran an initial 50000 iterations (100 ns) and checked convergence using the all-atom peptide radius of gyration (Rg) computed by the Python package *MDTraj v1.9.8*.[12]

To assess the REMD convergence, we adopted the temperature and radius of gyration analyses outlined in.[13] For temperature, we checked whether the replicas have sufficiently explored the various temperature states. Fig. S12 displays temperature trajectories of the first and last replicas in each simulation. Fig. S13 shows the percentage of time each replica spends in different temperature states. The data indicate that the replicas comprehensively traverse the entire temperature ranges, spending relatively even time across the states.

For radius of gyration (Rg), we plot Rg distributions from two distinct time intervals in Fig. S14. Simulations of PDB 6uf7, LowEnergy 19384, and the random sequences of 6uf7, 169032, and 31759, achieve observable Rg overlaps before 100 ns. For other simulations, the extension of runtime leads to Rg overlaps later on, with exceptions of Design 136805, Design 3114, LowEnergy 83218, LowEnergy 19384, LowEnergy 31759, and random sequence 31759. For these six cases, we analyze their average RMSDs over time and observe convergence, as illustrated in Fig. S15. Consequently, we conclude our REMD simulations as converged.

Figure S6: **CyclicChamp design results.** We list the number of structures obtained in each step of our CyclicChamp design workflow. For relaxed backbones and designed sequences, we show the number of structures with energies below the threshold (in kcal/mol). For high $P_{Near}$ candidates, we show the number of designs having $P_{Near} > 0.9$ out of the total validated designs, with the highest $P_{Near}$ score written below.

| n | Initial Points | Backbones | Clusters | Low-Energy Clusters | Designed Sequences | High Pnear (>0.9) Candidates | | | |
|---|---|---|---|---|---|---|---|---|---|
| | | | | | | *simple_cycpep_predict* | Rama-stability filtering | ClusterGen | Reshape |
| 7 | 39,996 | 5,277,863 | 12,196 | 6,981 (<8) | 513 (<−8) | 38/513 (0.981) | 77/513 (0.986) | – | – |
| 15 | 100,000 | 640,477 | 192,864 | 74,626 (<0) | 1,103 (<−30) | 3/75 (0.967) | – | 9/75 (0.958) | – |
| 20 | 100,000 | 983,567 | 157,217 | 81,718 (<0) | 97 (<−40) | 0/22 (0.172) | – | 0/69 (0.897) | 22/69 (0.982) |
| 24 | 100,000 | 186,483 | 42,758 | 26,537 (<0) | 105 (<−45) | 0/11 (0.042) | – | 0/82 (0.896) | 14/82 (0.991) |

Figure S7: **D-amino acid counts in CyclicChamp designs.** For each macrocycle size, we show the distribution of designs that have various number of D-amino acids. All designs have mixed chirality.

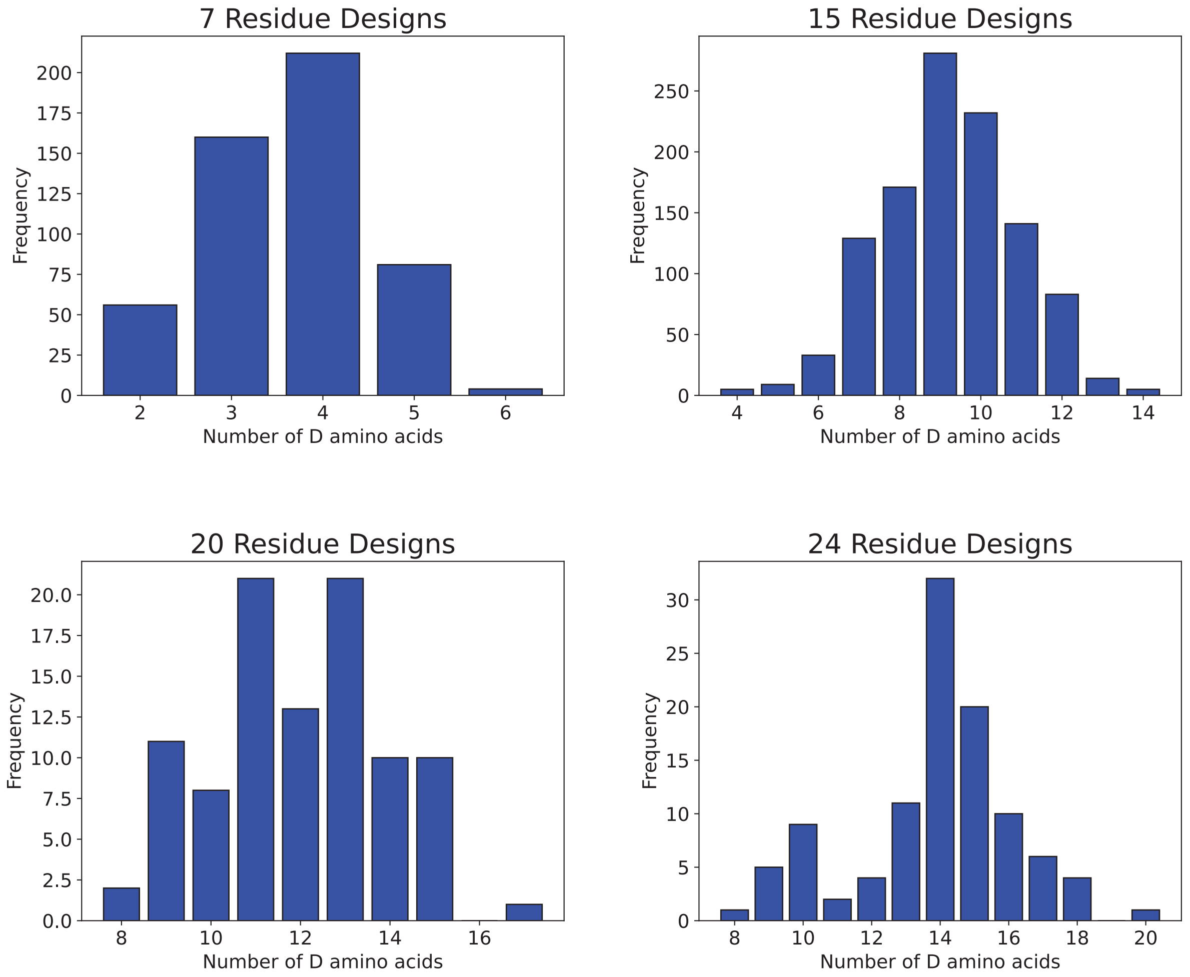

Table S2: Amino acid sequences of designs having high $P_{Near}$ values.

| **Top 7 Residue Designs** | |
|---|---|
| **Name** | **Sequence** |
| Design5191 | PRO-DPRO-PRO-DLYS-GLU-DASP-DHIS |
| Design10566 | VAL-DASN-GLU-DGLU-DPRO-PRO-DLYS |
| Design5790 | LYS-THR-DILE-DPRO-PRO-DVAL-ASP |
| Design9104 | PRO-DPRO-GLU-DGLN-GLU-ASN-DLYS |
| Design2787 | ASP-LYS-THR-DILE-DPRO-PRO-DVAL |
| Design2198 | ASN-DLYS-SER-GLU-DPRO-DTYR-DPRO |
| Design10632 | PRO-DPRO-LYS-DGLN-ASP-GLU-DGLN |
| Design372 | ASP-GLU-DPRO-DTYR-DPRO-ASN-DLYS |
| Design5982 | PRO-PRO-DPRO-GLU-DALA-LYS-DASN |
| Design9541 | DASN-PRO-DPRO-GLU-DLYS-LEU-DGLU |
| Design11103 | DASN-DPRO-ASN-DLYS-TYR-DPRO-GLU |
| Design1058 | GLU-DHIS-PRO-DPRO-DASP-DASP-DLYS |
| Design3860 | DSER-DLYS-PRO-DASN-DPRO-GLU-DVAL |
| Design8906 | PRO-ARG-DPRO-DARG-ASN-DASN-ASP |
| Design8145 | VAL-DASN-DLYS-DTHR-GLU-PRO-DPRO |
| Design2547 | VAL-DASN-DLYS-DSER-GLU-PRO-DPRO |
| Design5063 | GLU-DVAL-DPRO-DASN-PRO-DLYS-DPRO |
| Design9191 | PRO-ASN-DASN-GLU-DPRO-DLYS-SER |
| Design7437 | VAL-ASN-DASN-GLU-GLU-DPRO-DLYS |
| Design4816 | ASN-DSER-GLU-DPRO-LYS-PRO-THR |
| Design878 | DSER-GLU-DPRO-LYS-PRO-THR-ASN |
| Design9772 | GLU-DPRO-PRO-ARG-ASN-DTHR-SER |
| Design9974 | DGLU-PRO-LYS-DVAL-DASN-GLU-ASN |
| Design5863 | PRO-GLU-DSER-DPRO-DASN-ASN-DLYS |
| Design1828 | SER-PRO-ASN-DASN-GLU-DPRO-DLYS |
| Design1644 | DGLN-DPRO-PRO-THR-DASN-GLU-DARG |
| Design3836 | PRO-GLU-DASN-LYS-DASN-THR-GLU |
| Design3227 | PRO-THR-DASN-GLU-DARG-DGLN-DPRO |
| Design5621 | PRO-ASP-LYS-DASN-VAL-ALA-DPRO |
| Design7997 | GLU-PRO-GLU-DASN-LYS-DASN-THR |
| Design151 | PRO-ASN-GLU-DLYS-VAL-ALA-DPRO |
| Design12024 | DASN-ASN-DLYS-PRO-GLU-DSER-DPRO |

| Top 15 Residue Designs | |
|---|---|
| **Name** | **Sequence** |
| Design169032 | DTYR-DILE-DGLU-PRO-DVAL-ILE-PRO-SER-DSER-DGLU-PRO-TYR-DLYS-GLU-SER |
| Design3114 | ASP-GLU-SER-SER-DASN-DALA-DGLU-PRO-LYS-DSER-THR-DLYS-DPRO-ASN-DGLU |
| Design116427 | ASP-GLU-SER-DLEU-TYR-DALA-PRO-DPRO-TRP-ALA-ASN-DASP-DPRO-DARG-LEU |
| Design120532 | LYS-ASN-VAL-DGLN-DSER-ASP-LYS-DARG-DVAL-DPRO-DPRO-DLEU-DPRO-DASP-DTYR |
| Design16897 | THR-DSER-LYS-DLYS-PRO-DASP-DGLU-DARG-DALA-DASN-ASN-DSER-GLU-PRO-DSER |
| Design32090 | GLU-ALA-PRO-DARG-DPRO-DLEU-GLU-DPRO-TYR-DGLN-DTYR-DSER-DASN-ASN-DSER |
| Design17434 | DLEU-DARG-GLU-DTYR-DLEU-DASP-DGLU-PRO-PRO-GLU-LYS-ALA-DGLN-DASN-DASP |
| Design185692 | GLU-PRO-LYS-DTHR-DSER-DGLN-DALA-DARG-DLYS-DPRO-PRO-ASP-ASP-ARG-DALA |
| Design136805 | PRO-PRO-GLU-DSER-DALA-GLN-DSER-DSER-DGLU-DSER-PRO-DPRO-ALA-DLYS-DVAL |
| Design166081 | ALA-THR-ASN-DSER-DGLN-DLYS-DPRO-PRO-DASP-DALA-DTHR-DILE-THR-DLYS-ASP |
| Design167736 | PRO-DPRO-SER-DGLU-DPRO-DSER-DSER-DPRO-GLU-DALA-LYS-DALA-DPRO-ASN-DSER |
| Design2599 | GLU-GLU-DASN-GLU-DALA-SER-DTYR-DGLU-PRO-PRO-DTHR-DLYS-DGLU-DVAL-ASP |

| Top 20 Residue Designs | |
|---|---|
| **Name** | **Sequence** |
| Design24037 | DLYS-DSER-DSER-THR-ASN-DGLU-DASN-GLU-DALA-ARG-DGLU-DPRO-PRO-DILE-LEU-DGLU-DSER-DGLU-ASN-DALA |
| Design80837 | PRO-DPRO-DPRO-ALA-ALA-ASN-GLU-SER-DSER-THR-DASP-DGLU-DTHR-DALA-ASN-DASN-ASN-DASN-LYS-DGLU |
| Design25226 | DLYS-HIS-DLYS-PRO-SER-DLYS-ASP-LEU-LYS-DGLU-DALA-DGLN-PRO-TYR-DSER-DSER-DASN-ASP-ALA-LYS |
| Design119115 | DASN-PRO-DGLN-DALA-LYS-DALA-ASN-DTHR-TYR-ASP-ASP-ARG-DARG-SER-DGLU-DALA-GLU-DSER-GLN-DPRO |
| Design22588 | THR-DGLU-PRO-DPRO-DALA-GLU-ASP-GLU-DALA-DARG-GLU-SER-LEU-DALA-DLYS-PRO-DVAL-DHIS-DLYS-DLEU |

| | |
|---|---|
| Design74102 | DALA-DLYS-DHIS-DPRO-ASN-GLU-DASN-THR-SER-GLU-ALA-DGLN-DGLU-DPRO-DARG-ASN-DGLU-PRO-ALA-DASP |
| Design143166 | DASN-ASP-LEU-SER-DALA-DARG-DPRO-PRO-DGLU-GLN-DPRO-DTHR-LYS-ASP-GLU-DTHR-GLN-LYS-DSER-SER |
| Design45902 | DLYS-HIS-DLYS-PRO-THR-DLYS-ASP-PHE-LYS-DGLU-DALA-DGLN-PRO-TYR-DSER-DSER-DASN-ASP-ALA-LYS |
| Design26034 | DLYS-HIS-DLYS-PRO-THR-DGLU-LYS-PHE-LYS-DGLU-DALA-DGLN-PRO-TYR-DSER-DSER-DASN-ASP-ALA-LYS |
| Design1665 | DALA-ASP-ASP-ALA-PRO-DASP-DALA-DSER-ALA-ASP-LYS-DTHR-DMET-DPRO-DPRO-LYS-DASP-DGLU-DARG-THR |
| Design111 | TYR-DHIS-DGLU-DHIS-DASP-DGLU-DSER-THR-DTYR-LYS-DTHR-DGLU-PRO-DTHR-DPRO-ALA-DGLN-GLU-DTHR-DASN |
| Design83218 | ASP-ARG-DASN-DLYS-PRO-DLYS-DASP-DPRO-DVAL-TYR-DGLU-PRO-DPRO-DASN-DALA-DASN-DLYS-DSER-TYR-ASN |
| Design107505 | DLYS-DPRO-ASN-DTYR-ASN-DPRO-DLYS-DLEU-SER-DGLU-PRO-DASN-DTHR-DASN-GLU-DPRO-GLU-DARG-DTHR-DALA |
| Design59754 | ALA-DTHR-DTHR-DGLU-DTHR-DSER-DSER-DTYR-PRO-PRO-ASP-GLU-DLYS-THR-DTHR-DSER-DASN-GLU-DVAL-DHIS |
| Design12800 | DPRO-PRO-DARG-DPHE-DASN-ASN-DGLN-ILE-PRO-DASN-SER-DLYS-DPRO-DASP-DLEU-DGLN-ASP-LEU-SER-DGLU |
| Design15036 | TYR-PRO-DGLU-DALA-DSER-SER-LYS-DASP-DASP-DALA-DGLU-ASP-PRO-DLYS-DALA-DARG-DLYS-DLEU-GLN-DSER |
| Design27893 | DGLN-SER-DPRO-SER-ASP-GLN-SER-PRO-LYS-DLYS-ASN-DASP-DGLU-PRO-LEU-DSER-DASP-DGLU-DTYR-DASN |
| Design68384 | SER-GLU-DASN-ALA-LYS-DASP-DLEU-DGLU-PRO-DILE-DALA-DPRO-DASN-TYR-DPRO-DTHR-DPRO-LYS-DALA-PRO |
| Design23193 | SER-GLU-ALA-DLYS-DASN-DALA-DPRO-PRO-DSER-DPRO-DSER-DASP-DPRO-DSER-LYS-DASN-DGLU-PRO-VAL-DTYR |
| Design35869 | PRO-ASP-ALA-ASN-DGLU-ASP-MET-DALA-ALA-DSER-ILE-LYS-DGLU-GLU-DGLU-ASN-DSER-DARG-DPRO-DGLU |
| Design101360 | ASP-LYS-DSER-DLEU-LYS-DLYS-DASP-DASP-DVAL-DASP-DALA-DASP-DGLU-PRO-DVAL-ALA-LYS-PRO-ASN-DGLU |
| Design17670 | THR-SER-PRO-DALA-LYS-DASP-DLEU-GLU-DLYS-DASN-THR-DLYS-DASP-DALA-DPRO-DPRO-DARG-TYR-DALA-DGLU |

| Top 24 Residue Designs | |
|---|---|
| **Name** | **Sequence** |
| Design759 | LYS-DASP-DASP-DLEU-DLYS-ASN-DLEU-DTHR-DGLU-DPRO-LEU-DASN-PRO-DILE-TYR-SER-ASN-DGLN-DALA-ALA-DLYS-DASN-DALA-GLU |
| Design10052 | SER-DGLU-ALA-DPRO-DTHR-DALA-DALA-ALA-DPRO-PRO-DILE-GLU-LEU-DSER-DTHR-ASP-ALA-THR-ASN-DALA-ASN-DASP-DLYS-DASN |
| Design32190 | PRO-ASP-LYS-DASP-DALA-PRO-DASN-DTHR-DGLU-DPRO-DHIS-DGLU-DTYR-DGLU-DPRO-DLYS-PRO-DASN-ALA-DTYR-DGLU-DGLU-ALA-DLYS |
| Design21660 | THR-ALA-DALA-ALA-DALA-PRO-DGLU-DASN-DHIS-LYS-DPRO-DSER-DGLN-DPRO-DALA-GLU-DASN-GLU-DGLU-DTHR-THR-DASP-DLEU-DLYS |
| Design36199 | ALA-SER-DVAL-DSER-DGLU-THR-DVAL-DGLU-DPRO-DASP-DLYS-DPRO-GLU-DLYS-DALA-DTHR-PRO-DALA-DALA-ASN-DTHR-DASP-DALA-DSER |
| Design16647 | ASN-DGLN-SER-SER-DTHR-TYR-DALA-THR-DASP-DPRO-DTHR-PRO-DLEU-DALA-DTHR-DGLU-PRO-DASP-DARG-DARG-ALA-DGLN-ASP-ALA |
| Design37605 | DASP-DLYS-DARG-DALA-DARG-DASN-DALA-DGLU-DTHR-DGLU-SER-DALA-ASP-LEU-PRO-DASN-DSER-GLU-LYS-DASP-DLYS-PRO-THR-ALA |
| Design31759 | SER-DTHR-SER-SER-DGLU-LYS-DTYR-THR-DASP-DTYR-DARG-PRO-DLYS-DALA-DPRO-DPRO-PRO-DASP-DGLN-DALA-ALA-DGLN-ASP-MET |
| Design25212 | SER-GLU-THR-DGLU-SER-DHIS-PRO-DSER-SER-ARG-DASN-ASP-ASP-GLN-DASN-LEU-GLN-DALA-LEU-DVAL-PRO-DALA-THR-DASN |
| Design15225 | ALA-ASN-DTHR-ALA-DTHR-DSER-THR-DLYS-TYR-DASN-ASN-DASP-DALA-DGLU-DVAL-DTHR-PRO-DSER-GLU-PRO-LYS-DGLU-DASN-ASN |
| Design2901 | DSER-LYS-DGLU-PRO-DTYR-DASN-DVAL-DGLU-ASN-SER-DGLU-DGLN-DSER-DLEU-ARG-ASP-GLU-DGLU-LYS-DALA-DVAL-PRO-DASN-SER |
| Design18496 | THR-ASN-DLEU-DALA-DGLN-ALA-ARG-DTHR-DTHR-DASP-DILE-DTYR-DLYS-DPRO-PRO-THR-DARG-DPRO-SER-DALA-ASP-GLN-GLU-DLYS |
| Design20199 | DPRO-ARG-DGLU-DGLN-DTHR-TYR-DGLU-DASP-DVAL-THR-THR-SER-DGLU-DPRO-GLU-DLYS-PRO-DLYS-DALA-DALA-THR-DSER-DLEU-DLYS |
| Design27953 | THR-ASP-HIS-DSER-DGLU-PRO-SER-DLYS-SER-DALA-DASP-DGLU-DARG-DLYS-DTYR-PHE-ASP-DHIS-DARG-PRO-DGLN-DGLU-LYS-DPRO |
| Design19384 | DTYR-LEU-PRO-GLU-LEU-DSER-ALA-GLN-DGLU-PRO-ALA-DTHR-ALA-DASP-DLYS-DARG-DALA-DGLU-DPRO-DARG-SER-DSER-THR-DTYR |
| Design21698 | DSER-THR-PRO-ALA-LYS-SER-ASN-DVAL-PRO-LEU-DASP-DARG-DALA-DLYS-HIS-ASP-ASP-LYS-DARG-ASP-PRO-DGLN-DALA-ASN |

Figure S8: **Top 20-residue designs shown in sphere mode.** Prolines are colored in purple, and hydrophobic amino acids (ALA, ILE, LEU, VAL, MET, PHE) colored in orange.

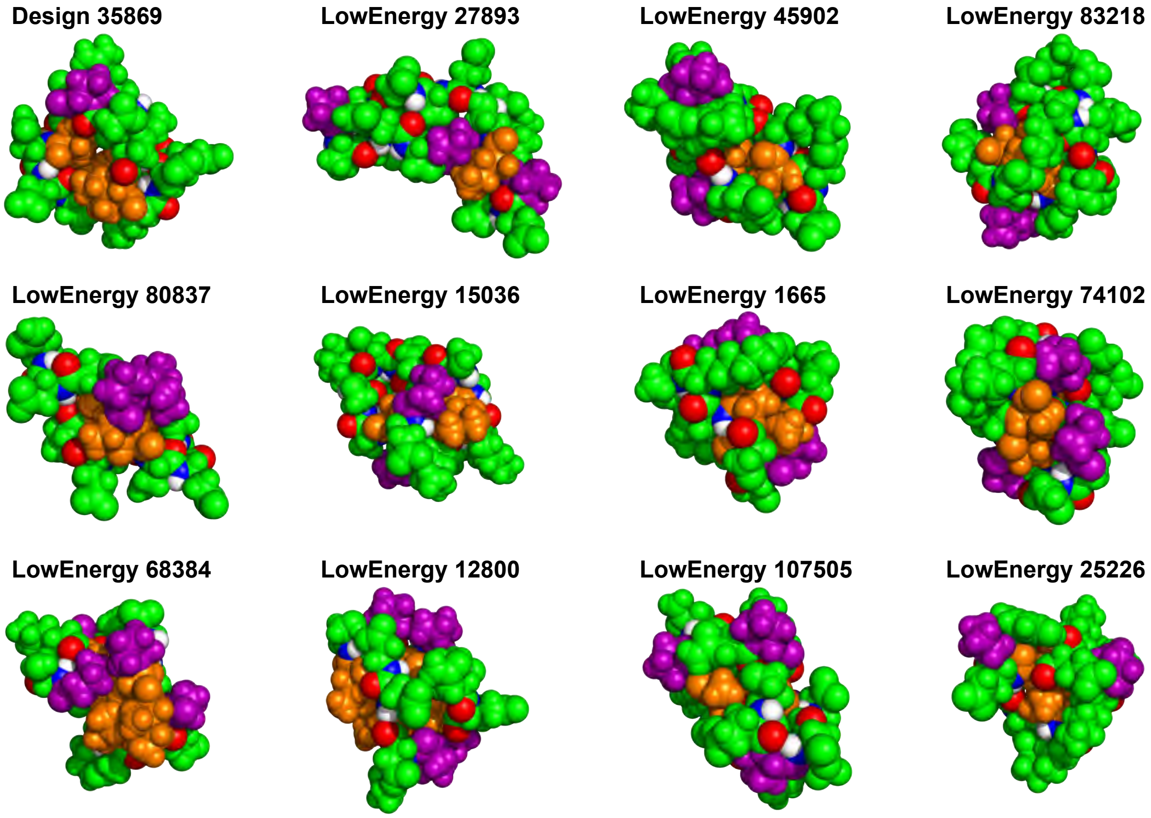

Figure S9: **Top 24-residue designs shown in sphere mode.** Prolines are colored in purple, and hydrophobic amino acids (ALA, ILE, LEU, VAL, MET, PHE) colored in orange.

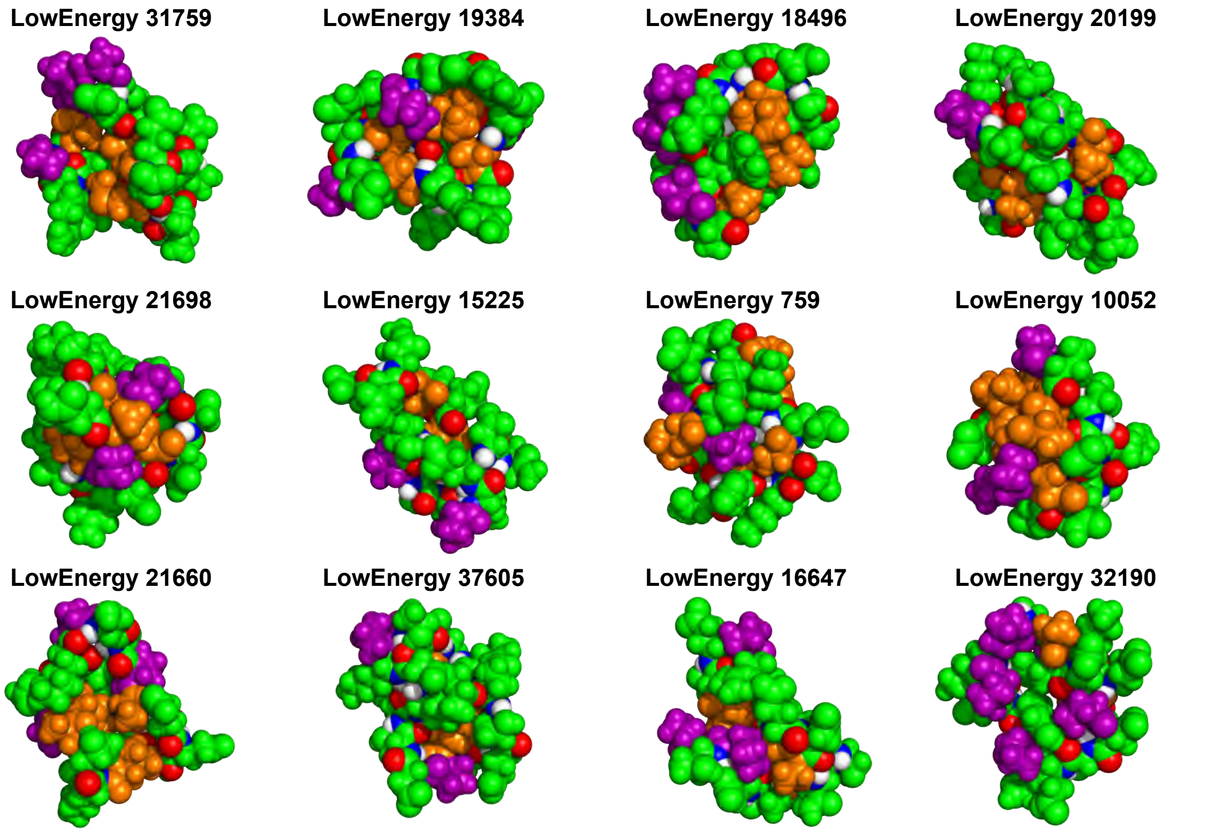

Figure S10: **Molecular dynamics simulation stable trajectories.** RMSDs are calculated between the backbone $C^{\alpha}$ atoms of the trajectory frames and our designed structures.

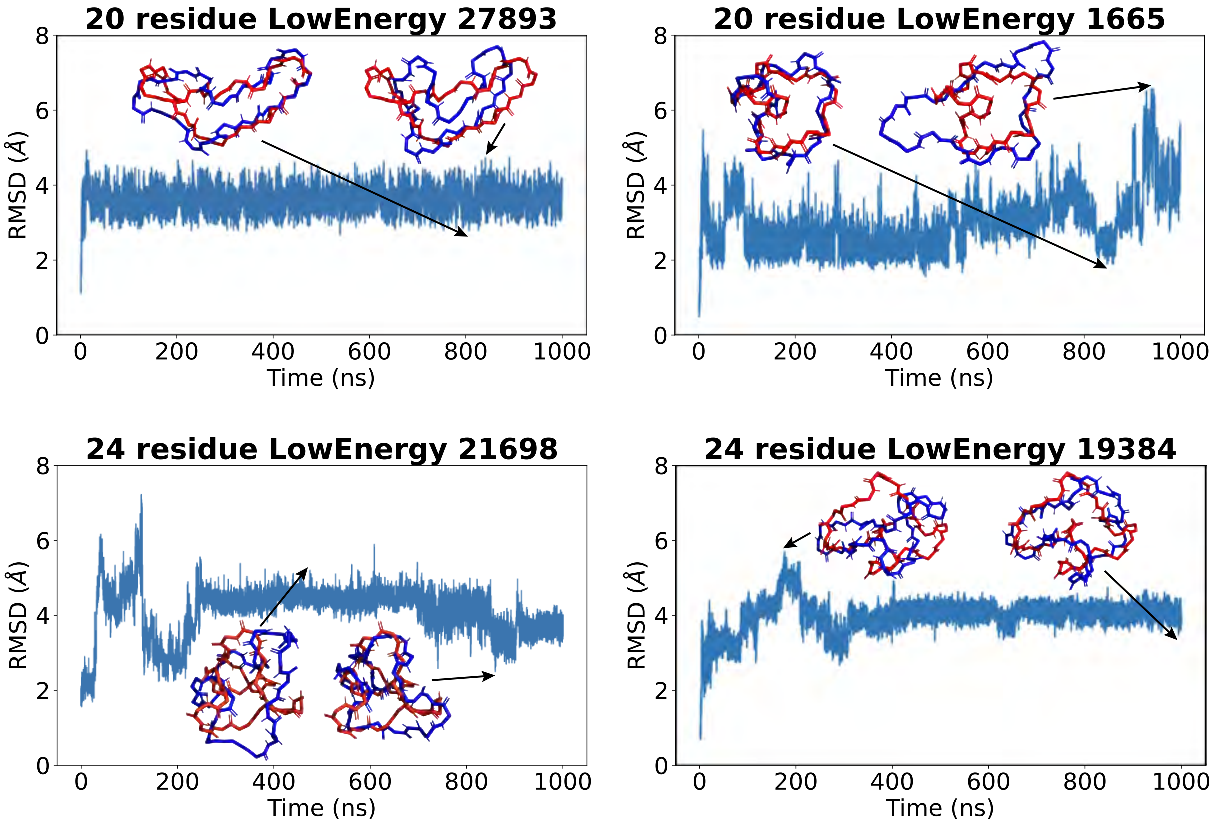

Figure S11: **Molecular dynamics simulation unstable trajectories.**

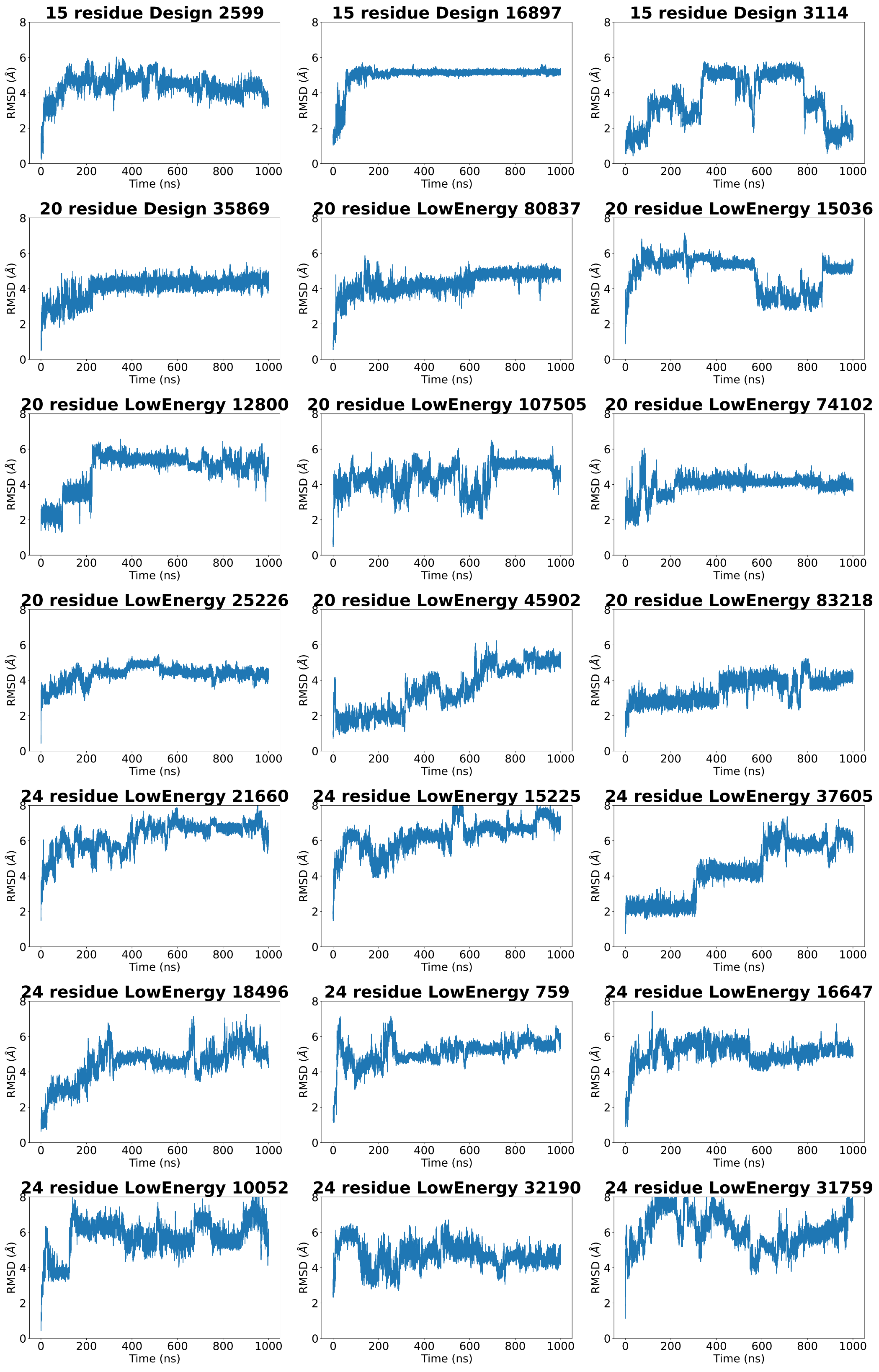

Figure S12: **REMD temperature plots.** For each simulation, temperatures of the first and the last replicas are drawn.

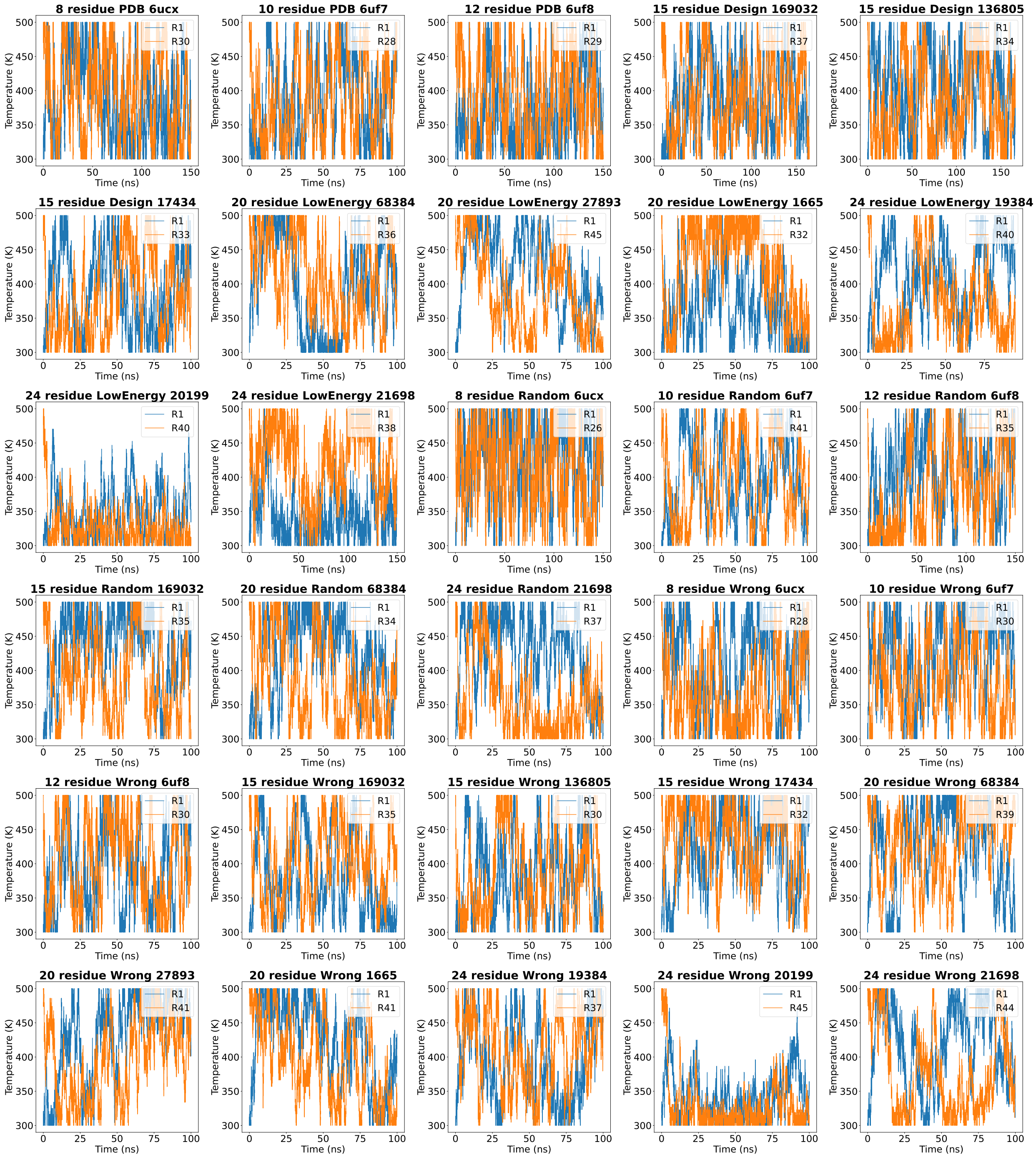

Figure S13: **REMD temperature dwell time plots.** The percentages of simulation time spent on each temperature are drawn for all replicas, with the standard deviations shown as error bars.

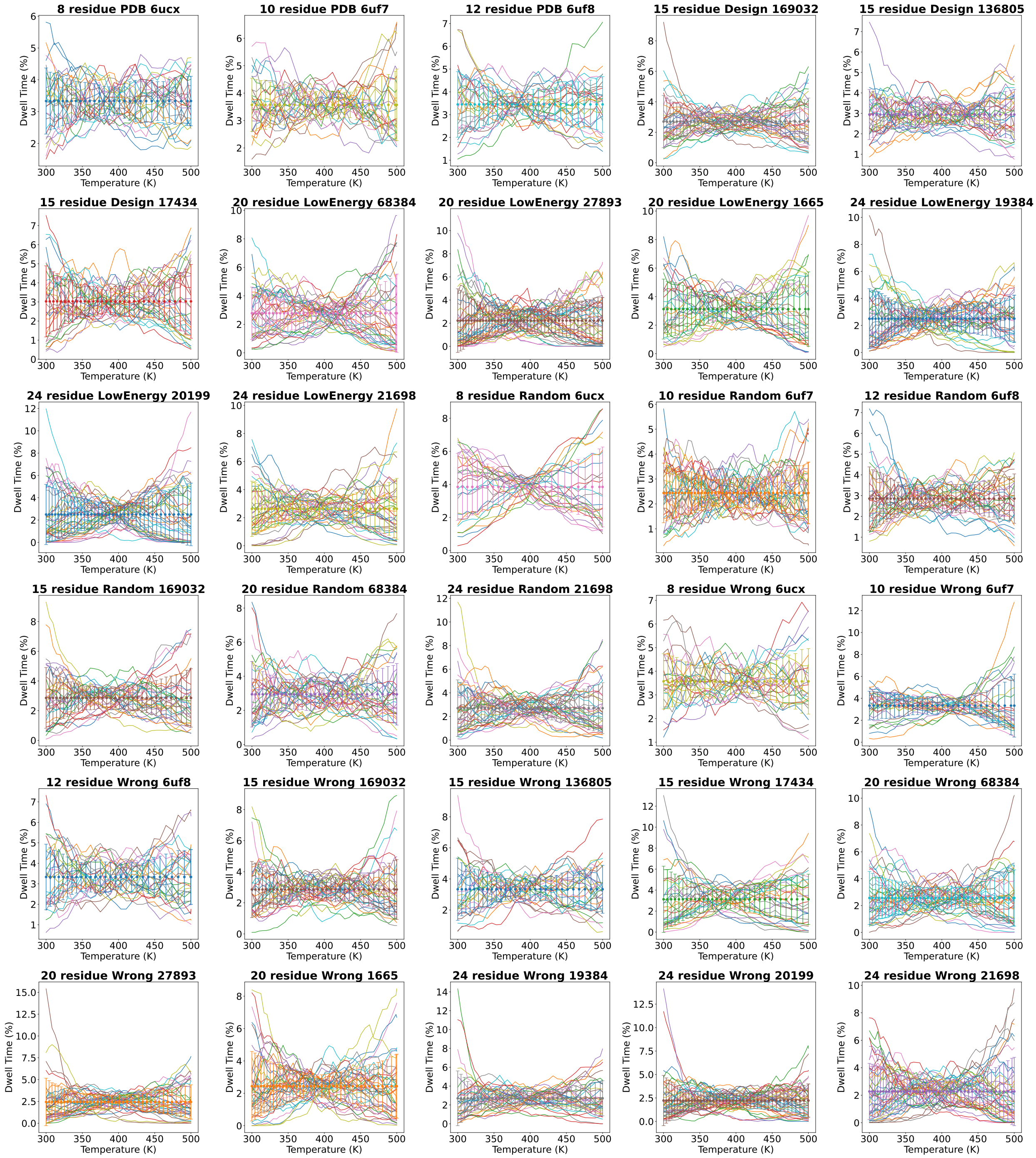

Figure S14: **REMD convergence check of Rg.** Distributions of all-atom peptide radii of gyration are plotted for two different time intervals.

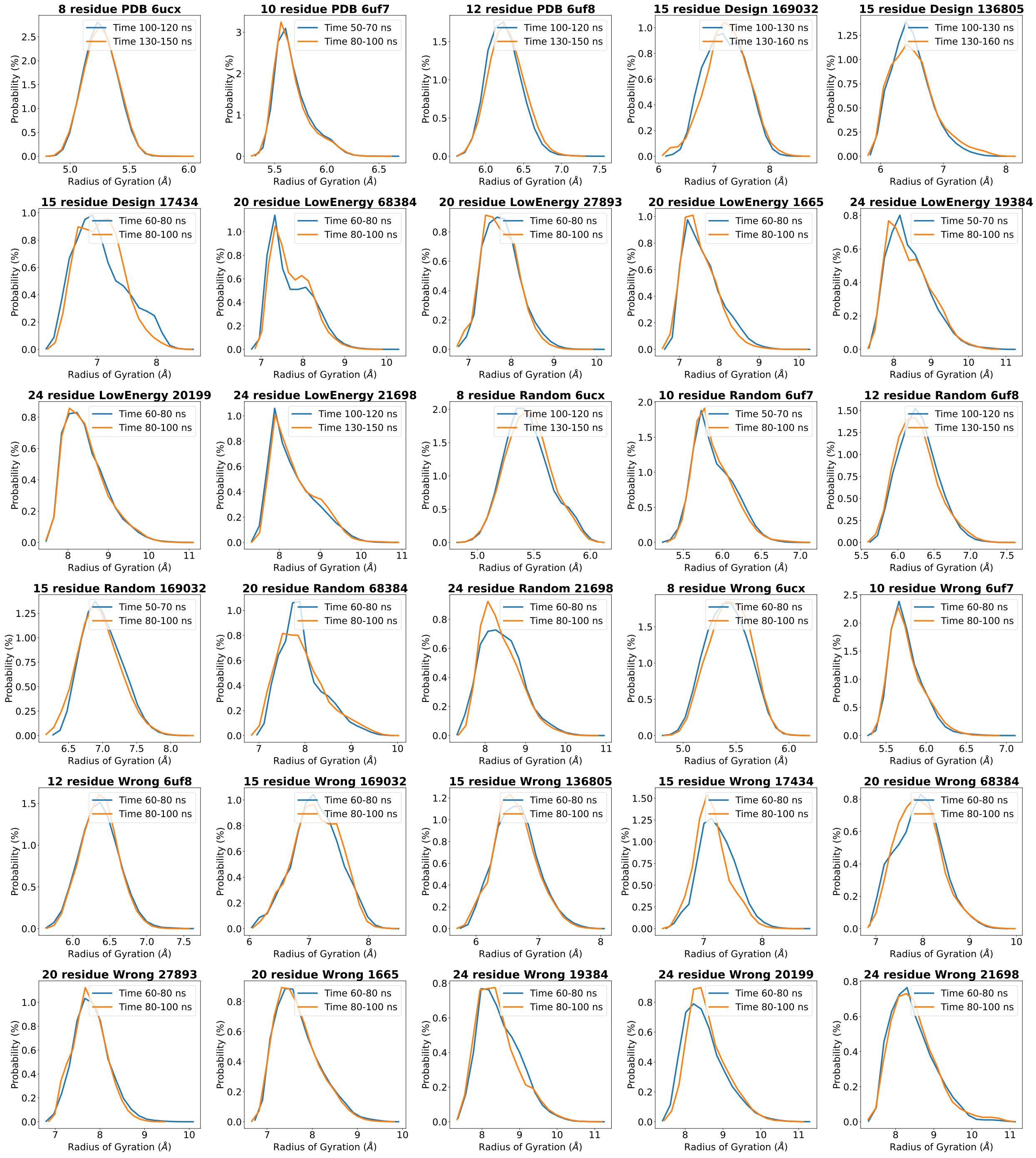

Figure S15: **REMD convergence check of RMSD.** For simulations that do not show a good overlap of Rg distributions from the two time intervals, we plot the average $C^{\alpha}$-atom RMSDs between trajectory frames sampled at temperature state 300 K and our designed structure over simulation time.

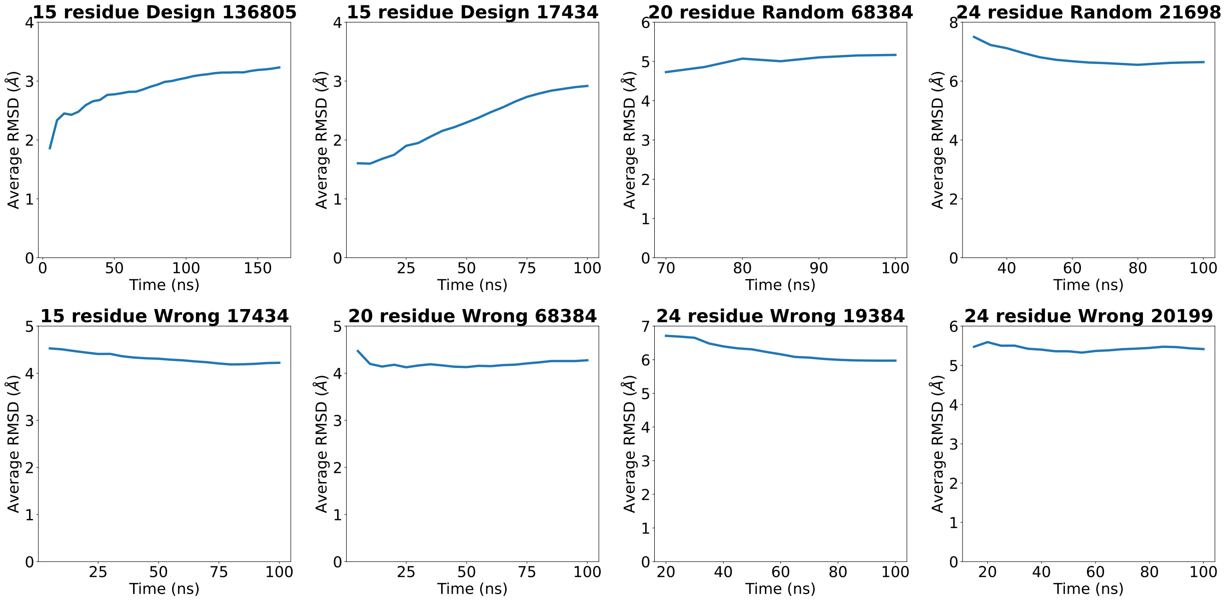

Figure S16: **REMD free energy surfaces with negative controls.** (a) As a negative control, for each size, we selected one design and randomly permuteed its sequence. The FES comparisons are shown. (b) For our 15-24 residue designs, we re-ran the REMD simulation from a distinct starting structure (colored in green) chosen from the $P_{Near}$ landscape, which has low energy and high RMSD from the design (2.1-6.8 Å).

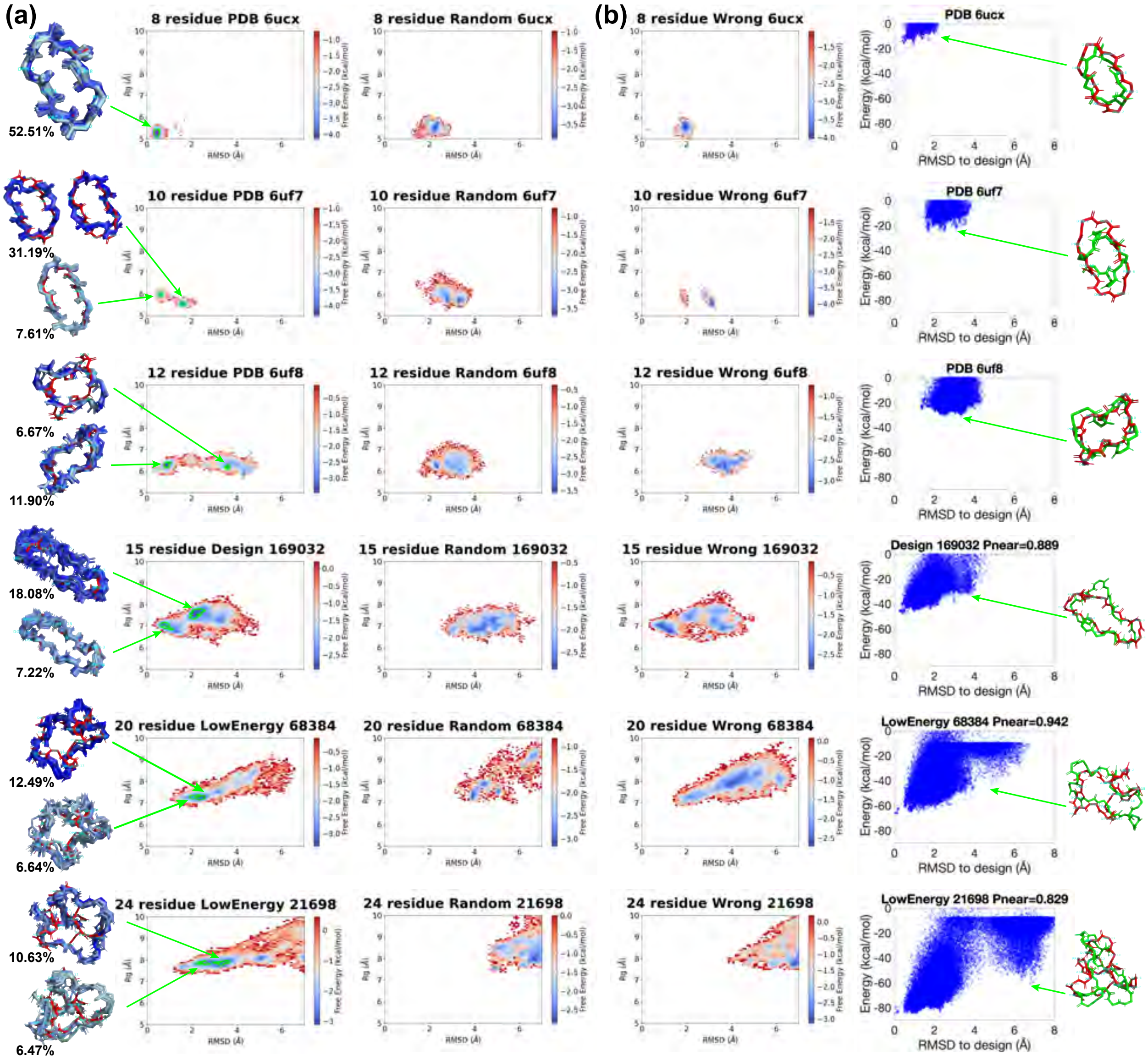

Figure S17: **Other REMD free energy surfaces.** (a) Representative structures are shown for the energy basins. (b) REMD simulations re-run from distinct starting structures (colored in green) chosen from the $P_{Near}$ landscapes, which have low energies and high RMSDs from the designs (3.8-6.8 Å).

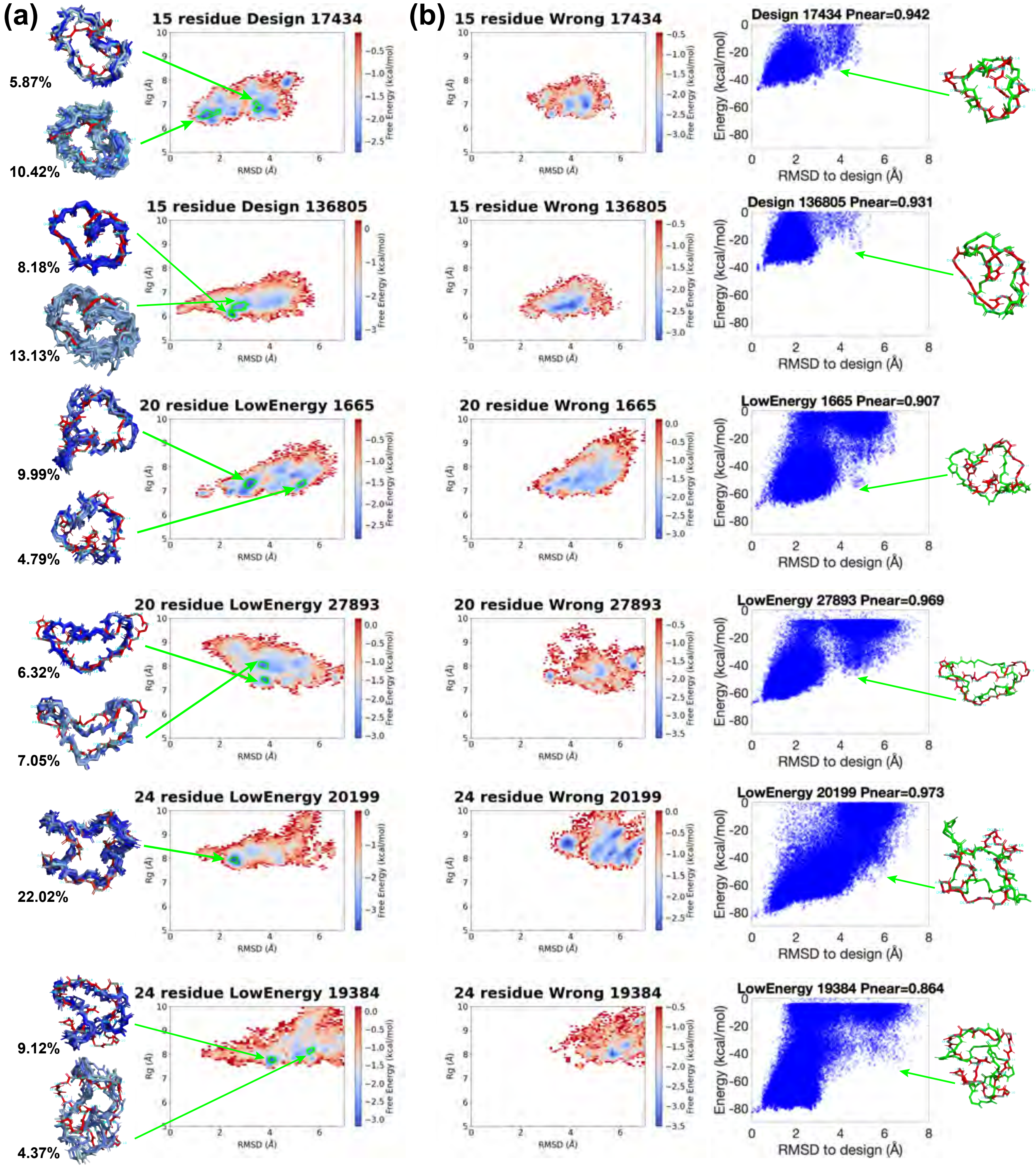

Figure S18: **Energy landscape predictions of available macrocycle structures deposited in the PDB.** In each landscape, the predicted low-energy cluster centers are marked in red. Rosetta energy scores of the PDB structures are drawn as dash lines for reference.

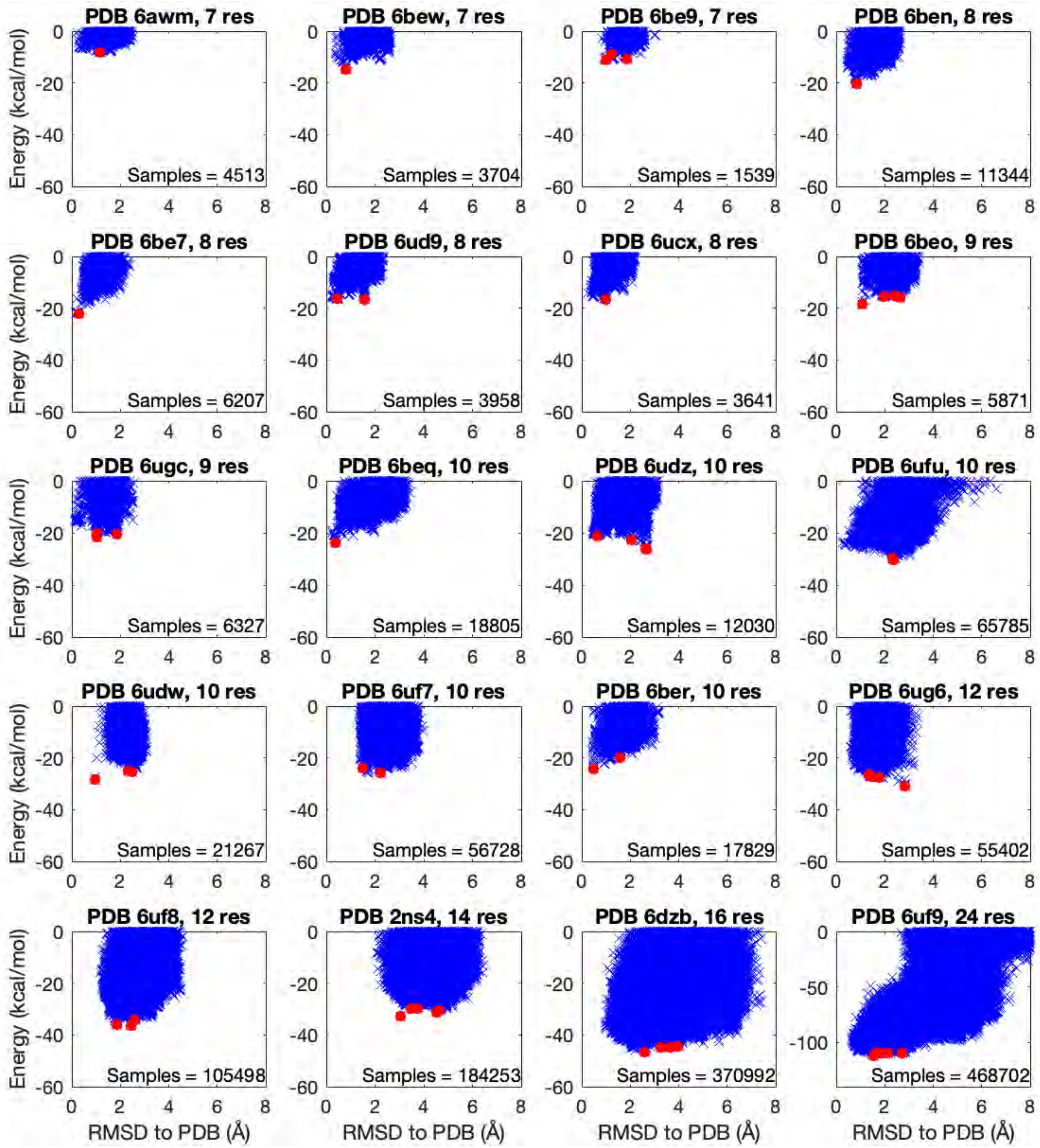

Figure S19: **Other structure predictions for available macrocycle structures deposited in the PDB.** Predicted low-energy cluster centers (green) are aligned to the PDB structures (orange), with RMSDs shown. PDB structures from the 2017[14] and 2020[15] Rosetta paper are labeled in blue and black, respectively, and all other structures in pink.

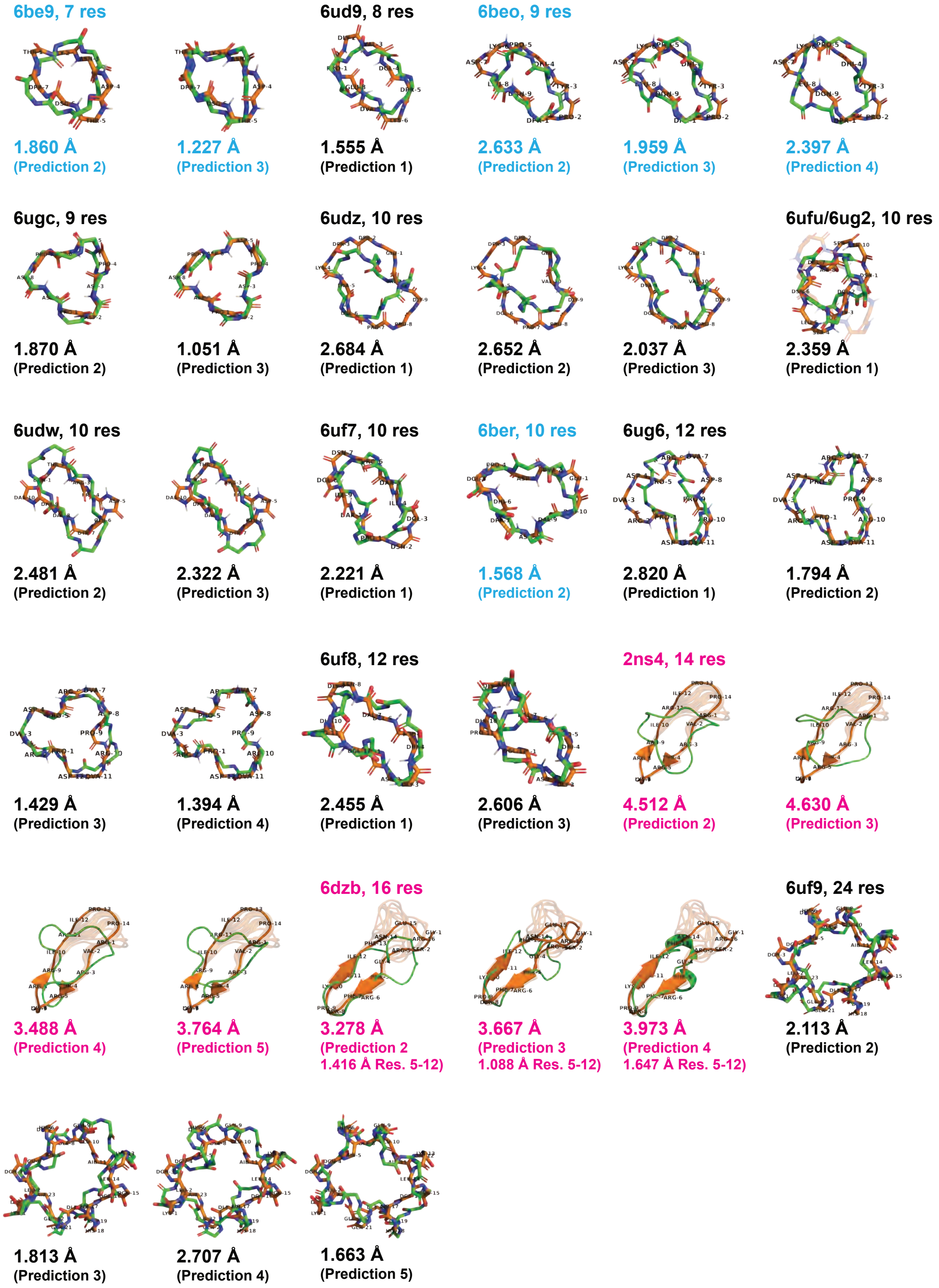
